# Identification of Host Restriction Factors Critical for Recombinant AAV Transduction of Polarized Human Airway Epithelium

**DOI:** 10.1101/2023.09.27.559795

**Authors:** Siyuan Hao, Xiujuan Zhang, Kang Ning, Zehua Feng, Soo Yeun Park, Cagla Aksu Kuz, Shane McFarlin, Donovan Richart, Fang Cheng, Elizabeth Yan Zhang, Aaron Zhang-Chen, Ziying Yan, Jianming Qiu

## Abstract

Recombinant (r)AAV2.5T was selected from the directed evolution of an AAV capsid library in human airway epithelium (HAE). The capsid gene of rAAV2.5T is a chimera of the N-terminal unique coding sequence of AAV2 VP1 unique (VP1u) and the VP2- and VP3-coding sequence of AAV5 with a single amino acid mutation of A581T. We conducted two rounds of genome wide CRISPR gRNA library screening for host factors limiting rAAV2.5T transduction in HeLa S3 cells. The screen identified several genes that are critical for rAAV2.5T transduction in HeLa S3 cells, including previously reported genes *KIAA0319L*, *TM9SF2*, *VPS51*, and *VPS54*, as well as a novel gene *WDR63*. We verified the role of KIAA0319L and WDR63 in rAAV2.5T transduction of polarized HAE by utilizing CRISPR gene knockouts. Although KIAA0319L, a proteinaceous receptor for multiple AAV serotypes, played an essential role in rAAV2.5T transduction of polarized HAE either from apical or basolateral side, our findings demonstrated that the internalization of rAAV2.5T was independent of KIAA0319L. Importantly, we confirmed WDR63 is an important player in rAAV2.5T transduction of HAE, while not being involved in vector internalization and nuclear entry. Furthermore, we identified that the basal stem cells of HAE can be significantly transduced by rAAV2.5T.

**Significance:** The essential steps of a successful gene delivery by rAAV include vector internalization, intracellular trafficking, nuclear import, uncoating, double-stranded (ds)DNA conversion, and transgene expression. rAAV2.5T has a chimeric capsid of AAV2 VP1u and AAV5 VP2 and VP3 with the mutation A581T. Our investigation revealed that KIAA0319L, the multiple AAV serotype receptor, is not essential for vector internalization but remains critical for efficient vector transduction to human airway epithelia. Additionally, we identified that a novel gene *WDR63*, whose cellular function is not well understood, plays an important role in vector transduction of human airway epithelia but not vector internalization and nuclear entry. Our study also discovered the substantial transduction potential of rAAV2.5T in basal stem cells of human airway epithelia, underscoring its utility in gene editing of human airways. Thus, the knowledge derived from this study holds promise for the advancement of gene therapy in the treatment of pulmonary genetic diseases.

## Introduction

Adeno-associated viruses (AAVs) are members of genus *Dependoparvovirus* of the *Parvoviridae* family (1). AAV is non-enveloped and has a single-stranded (ss)DNA genome of 4.7 kb. It is nonpathogenic to humans and replication-deficient *per se*. It requires a helper virus, e.g., adenovirus (Ad), to replicate. Recombinant (r)AAVs are considered a safe transduction vector for in vivo gene transfer in the application of human gene therapy (2,3). Since the late 1990s, rAAVs have been used in a number of clinical trials to treat a wide variety of monogenetic disorders across target organs, including the lung, eye, liver, and skeletal muscle (4–8). So far, the US Food and Drug Administration (FDA) has approved four rAAV-based gene therapy medicines, LUXTURNA, ZOLGENSMA, HEMGENIX, and ELEVIDYS for the treatment of Leber congenital amaurosis, spinal muscular atrophy, hemophilia B, and Duchenne muscular dystrophy, respectively.

One major hurdle in rAAV-based gene therapy is the requirement of a high vector dose. Although rAAVs have been extensively used in biomedical research and gene therapy, there is still limited understanding of how they enter cells and deliver their cargo to the nucleus. Additionally, the essential host factors that facilitate transgene expression vary depending on the specific AAV serotype and the type of cells being targeted (9). The cell and tissue tropism of an AAV is determined by its capsid structure, which is composed of viral proteins VP1, VP2, and VP3 at a molar ratio of ∼1:1:10 (10), and by its interaction with the specific surface receptors and host factors present on or within the target cells. In general, the cell attachment of most AAV serotypes is mediated by capsid-specific glycan moieties on the cell surface that has been studied extensively (11). After attachment, AAVs are internalized into the target cell through receptor-mediated endocytosis. Previously, a genetic screen identified a highly conserved AAV entry factor, KIAA0319L, as a cellular receptor for certain naturally occurring AAVs (AAV1, 3, 5, 6, 8, and 9) (12). However, it is important to note that AAV4-like AAVs (AAV4, AAV11, and AAV12) do not utilize KIAA0319L for vector transduction (13). KIAA0319L is classified as a type I transmembrane protein. The purified recombinant KIAA0319L or an anti-KIAA0319L antibody effectively inhibits transduction of rAAV1-3, 5, 6, 8, and 9, and knockout of *KIAA0319L* in in vitro cultured cell lines or mice resulted in resistance to the rAAV transduction (12). Although KIAA0319L was characterized as an N- and O-glycosylated cell membrane protein, the glycosylation does not contribute to AAV binding and transduction (14). It appears transiently on the cell surface but localizes primarily to the perinuclear trans-Golgi network (TGN). While the native role of KIAA0319L is presumed to involve acting as a recycling receptor for an unidentified ligand, it has been identified as a proteinaceous receptor for certain AAV serotypes. AAV has evolved to exploit the retrograde trafficking of KIAA0319L. However, the precise mechanism by which KIAA0319L facilitates rAAV transduction remains elusive, specifically, its involvement in AAV cell surface binding, AAV internalization, intracellular trafficking, potential interacting partners, and its recycling route from cell surface to the TGN.

Even though it remains unclear how AAV capsids differentially interact with host factors, AAV capsids can be engineered to modify their cell/tissue tropism by introducing specific mutations that enhance or alter their interactions with receptors and/or cellular factors relevant to the desired target cells or tissues. AAV2.5T is an airway tropic AAV variant identified from the directed evolution of an AAV capsid library in polarized human airway epithelia. The VP1 of AAV2.5T is a chimeric protein, comprised of the AAV2 VP1u region (amino acids (aa) 1-118) with the remaining amino acids (aa) 119-725 from AAV5 VP1, along with a key point mutation (A581T), and the VP2 and VP3 of AAV5 bearing the same mutation (15). rAAV2.5T vector demonstrated greater than 10-fold improvement over AAV2 and AAV5 in the transduction of human airway epithelia (15). It efficiently transduces cystic fibrosis (CF) human airway epithelia from the apical side. rAAV2.5T successfully delivered the expression of CF transmembrane conductance regulator minigene (CFTRΔR) with a 156-bp deletion in the regulatory (R) domain (16) to correct the deficient transepithelial transport of chloride anion in the polarized human airway epithelia derived from the CF lung donors (15,17).

Similar to its parent AAV5, AAV2.5T uses cell surface α2,3 N-linked sialic acid (SA) as the primary attachment receptor (18). The SA-binding sites of AAV5 were mapped to M569, A570, T571, G583, T584, Y585, N586, and L587, and the binding pocket has a contact distance of 2.4 to 3.6 Å to interact with SA (19). While the A581T mutation is close to the SA binding site, its impact on the SA binding is minimal due to the fact that the binding affinities of AAV5 and AAV2.5T to α2,3 N-linked SA remain similar (18), suggesting that both use the same mechanism to bind SA.

Understanding the important host factors that facilitate rAAV2.5T transduction in human airway epithelia is crucial to fully exploit the potential of this novel vector in CF gene therapy. In this study, we first undertook a genome wide CRISPR screen in Hela S3 cells. We found that gRNAs targeting the gene *KIAA0319L* had the highest numbers of reads from the analysis of the screen. Importantly, our study also identified that gene *WDR63* is important for AAV2.5T transduction of human airway epithelia. Interestingly, our vector internalization assays demonstrated that neither KIAA0319L nor WDR63 plays a role in the internalization of rAAV2.5T into the cells of human airway epithelia. Thus, our findings indicate that KIAA0319L may function differently in different types of cells and tissues for rAAV transduction.

## Results

### Genome wide CRISPR screen identifies novel restriction factors of rAAV2.5T transduction

The capsid of AAV2.5T differs from AAV5 by only a single amino acid (A581T) in VP2 and VP3, while the N-terminal of the VP1, which is replaced by the VP1u of AAV2, remains concealed inside the capsid. While AAV5 interacts with KIAA0319L to promote transduction in HeLa cells (14), it has been reported that rAAV2.5T transduction of HeLa cells or apical transduction of polarized human airway epithelia is independent of KIAA0319L (20), suggesting that rAAV2.5T and rAAV5 exploit different mechanisms for transduction. To explore the specific receptors and co-receptors involved in rAAV2.5T entry into human airway epithelia, we utilized a genome wide CRISPR pooled guide (g)RNA library for high throughput screening to identify the significant host factors that restricted rAAV2.5T transduction in HeLa S3 cells. To facilitate the screen, suspension HeLa S3 cells were transduced with a lentiviral vector to incorporate the expression of spCas-9. These spCas9-expressing HeLa S3 cells were transduced with the lentiviral-based pooled gRNA library (Brunello) for human gene knockout at a multiplicity of infection (MOI) of 0.25 transduction units per cell to ensure each cell was transduced by no more than one gRNA-expressing lentivirus. Following the selection of cells with puromycin at 2 µg/ml, the spCas9 and sgRNA-expressing Hela S3 cells were extracted for genomic DNA (gDNA), designated as gDNA^Ctrl^. For the screen, we transduced the spCas9 and sgRNA library-expressing HeLa S3 cells with mCherry-expressing rAAV2.5T (rAAV2.5T.F5tg83luc-CMVmCherry) at an MOI of 20,000 DNase resistant particles (DRP)/cell, by which a transduction efficiency of 96.4% (mCherry positive) was achieved. Using flow cytometry, we sorted ∼1 million mCherry-negative cells and amplified them for the second-round of transduction of rAAV2.5T at the same MOI. Again, ∼1 million mCherry negative cells were sorted, expanded, and underwent extraction of gDNA that was designated as gDNA^Screen^ (**Figure 1A**). Both the gDNA^Ctrl^ and gDNA^Screen^ were subjected to next generation sequencing (NGS) to probe the single gRNAs (sgRNAs) enriched in the transduction restricted cells. Analysis using the bioinformatics software MAGeCK highlighted the sgRNAs that were significantly enriched in the second round of sorted mCherry-negative cells, compared to unselected control cells (**Table S1** and **Figure S1**). The genes that have an enrichment score of >3.25 are shown in **Figure 1B**. Most notably, *KIAA0319L*, which was previously reported as a multi-serotype AAV receptor (12), was the top selected gene (**Figure 1B**). Gene ontology (GO) classification showed many potential rAAV2.5T host restriction factors identified in our screen, including TM9SF2, VPS51, and VPS54, which were previously identified as the host restriction factors for rAAV transduction (11).

**Figure 1.**
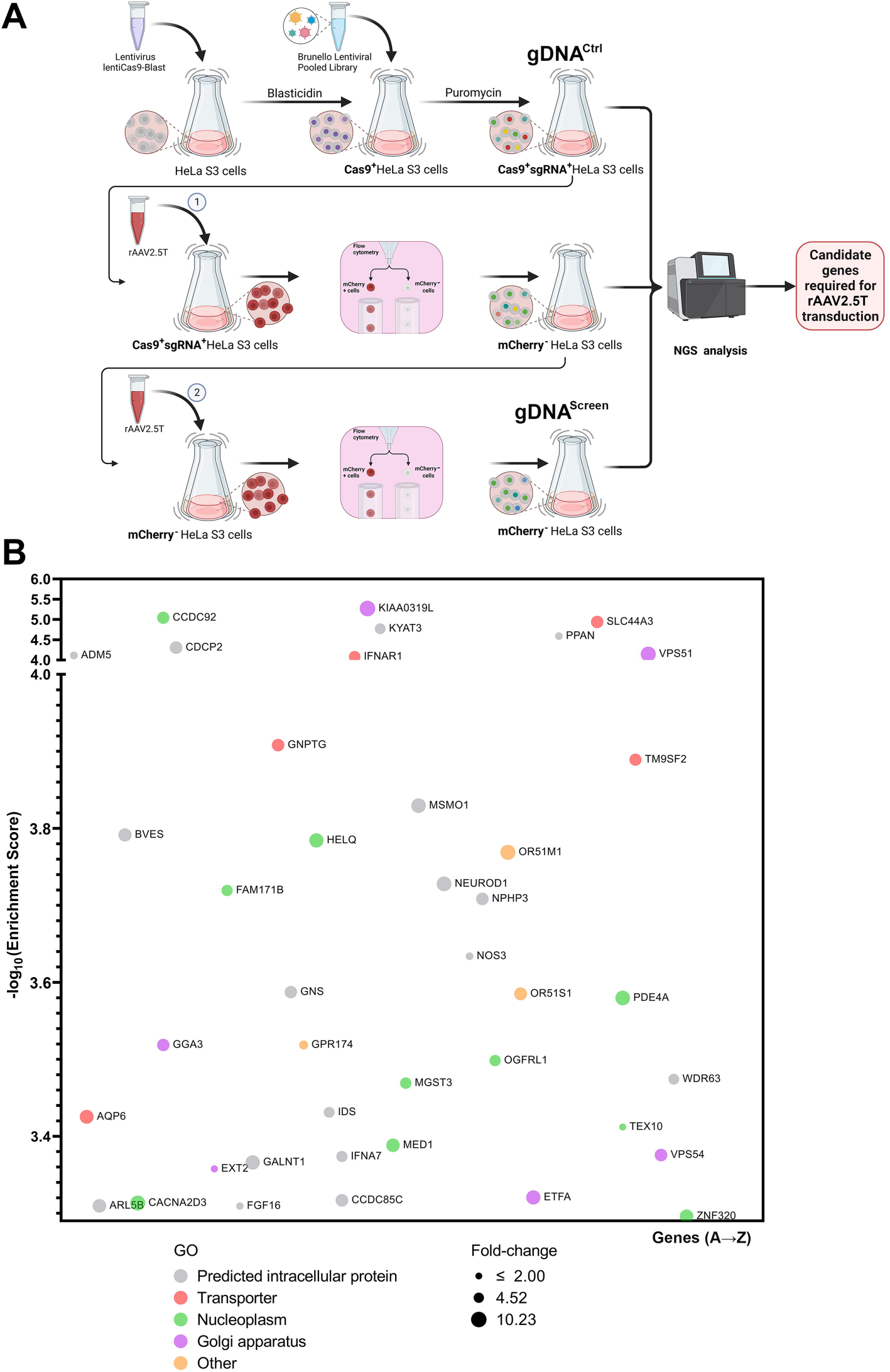
Genome-wide guide RNA (gRNA) library screen of restriction factors of rAAV2.5T transduction in HeLa S3 cells. **(A) A diagram of Genome-wide gRNA library screen**. HeLa S3 suspension cells were transduced with a spCas9 expressing lentivirus. Blasticidin-resistant spCas9-expressing cells were transduced with the Brunello lentiCRISPR gRNA lentiviral library. The library-selected HeLa S3 cells were expanded to ∼200 million, of which ∼100 million were extracted for genomic DNA (gDNA) as the non-selected control gDNA (gDNA^ctrl^), and another 100 million were transduced with mCherry-expressing rAAV2.5T. At 3 days post-transduction, ∼1 million mCherry negative cells were sorted and expanded. ∼100 million cells were transduced with rAAV2.5T, followed by sorting of the top 1% mCherry negative cells. The sorted mCherry negative cells were expanded to ∼100 million for extraction of gDNA^Screen^. The gDNA samples were subjected to NGS analysis and bioinformatics analysis. **(B) Hits of host genes from the second round of screen of mCherry negative cells.** This panel shows the entry screen hits from the second-round selection of mCherry-negative cells transduced with rAAV2.5T. The x-axis shows genes within the target of Brunello library and are grouped by gene ontology analysis. Each circle represents a gene with the size corresponding to fold change of sgRNA reads by comparison of gDNA^Screen^ to gDNA^ctrl^. The y-axis shows the enrichment score [-log_10_] of each gene based on MAGeCK analysis of the sgRNA reads in gDNA^Screen^ vs gDNA^ctrl^. Only genes that have an enrichment score of >3.25 are shown. The bubble colors indicate different gene ontology classifications as shown.

### HeLa cell screening identifies *KIAA0319L*, *TM9SF2*, and *WDR63* as important host genes

We selected the 27 top candidates with an enrichment score of >3.25 (**Figure 1B**) and available antibodies, to verify their roles in rAAV2.5T transduction in HeLa cells. We first used shRNA-expressing lentiviruses to selectively silence the expression of these candidate genes, respectively, in HeLa cells. The 27 candidate genes were silenced to different extents, as shown by Western blotting (**Figure S2**). Dual-reporter rAAV2.5 vector, rAAV2.5T.F5tg83luc-CMVmCherry, was used to transduce these individually gene-silenced HeLa cells at an MOI of 20,000. At three days post-infection, the transduction efficiencies were assessed for mCherry expression and luciferase activity using flow cytometry and firefly luciferase activity assays, respectively. After normalization of the mCherry expression with the scramble (non-target) shRNA control cells, the data showed that silencing of *KIAA0319L*, *KYAT3*, *TM9SF2*, *NPHP3*, *WDR63*, and *ZNF320* decreased mCherry expression by >50% (**Figure 2A**, columns in red). Silencing of *CCDC92*, *CDCP2*, *MSMO1*, *OR51M1*, *PDE4A*, and *GPR174* decreased mCherry expression by >25% but <50% (**Figure 2A**, columns in green). Similarly, we found that silencing of *KIAA0319L*, *TM9SF2*, *WDR63*, and *ZNF320* had a decrease in luciferase expression by >50% (**Figure 2B**, columns in red) and that silencing of *KYAT3*, *OR51M1*, *NPHP3*, and *PDE4A* decreased luciferase expression by >25% but <50% (**Figure 2B**, columns in green).

**Figure 2.**
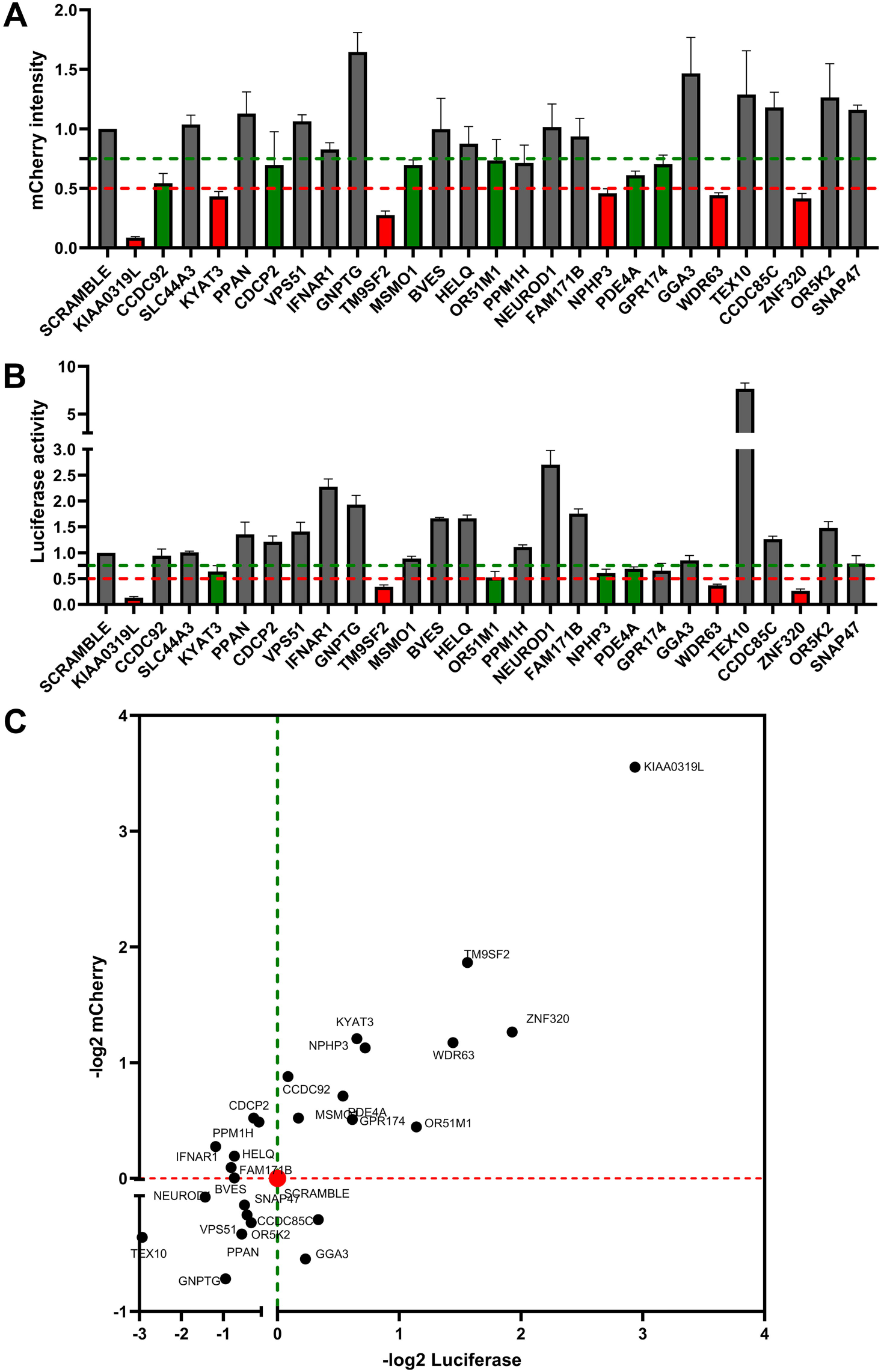
rAAV2.5T transduction in the top 28-ranking gene silenced HeLa cells. **(A&B) Relative expression of luciferase and mCherry in gene silenced cells.** HeLa cells seeded in 24-well plates were transduced with a shRNA-expressing lentiviral vector. After selection in puromycin, the cells were transduced with rAAV2.5T at an MOI of 20,000 DRP/cell. After 3 days post-transduction, mCherry expression was imaged under a fluorescent imager (A), and the luciferase activity was measured using a firefly luciferase detection reagent (B). All data had 3 repeats and were normalized to the non-target (NT) control. The green and red dashed lines indicated 75% and 50% of the NT control, respectively. Data shown are means with a standard deviation (SD) from three replicates. **(C) Ranks of the candidates in Quadrant-I.** X-axis shows −log_2_ luciferase intensity, and Y-axis shows −log_2_ mCherry intensity. All the cells silenced with gene names shown in Quadrant I had decreases in both luciferase and mCherry expressions.

We plotted relative expression values (-log_2_) of both luciferase and mCherry expressions (**Figure 2C**). The genes whose silencing decreased both expressions are shown in Quadrant I, with *KIAA0319L* being the highest and *TM9SF2* being the second-highest hit. Both have been shown to be essential entry factors of transduction in the whole genome gRNA library screens using rAAV2 (12,21). We finally narrowed the candidate restriction factor genes down to 11, i.e., *KIAA0319L*, *TM9SF2*, *ZNF320*, *WDR63*, *KYAT3*, *NPHP3*, *PDE4A*, *OR51M1*, *CCDC92*, *MSMO1*, and *GPR174*. Then, we looked back to our NGS data for these 11 genes (**Figure S3**), and found that most of the genes showed an average of >2,000-fold enrichment in sgRNA reads, compared to the unselected control, indicating that these candidates are highly likely to play a role in rAAV2.5T transduction.

Next, we examined the role of these 11 candidate genes in rAAV2.5T transduction in gene-knockout HeLa cells. To this end, HeLa cells were transduced with individual sgRNA-expressing lentiviral vector and underwent single cell expansion. Western blotting demonstrated drastic decrease in expression of all the selected candidate genes (**Figure 3A**). Then, each line of the gene-knockout cells was infected with rAAV2.5T. The results showed that *KIAA0319L* knockout decreased luciferase expression by >90%, and *TM9SF2* and *WDR63* knockout decreased luciferase expression by >60% (**Figure 3B**). Notably, *WDR63* knockout decreased luciferase expression to the same level as the *TM9SF2* knockout. In addition, *PDE4A*, and *GPR174* knockdown showed ∼50% decreases in the luciferase expression. However, *ZNF320*, *KYAT3*, and *CCDC92* gene knockout did not show any significant decrease in luciferase expression when compared to the non-target control (**Figure 3B**).

**Figure 3.**
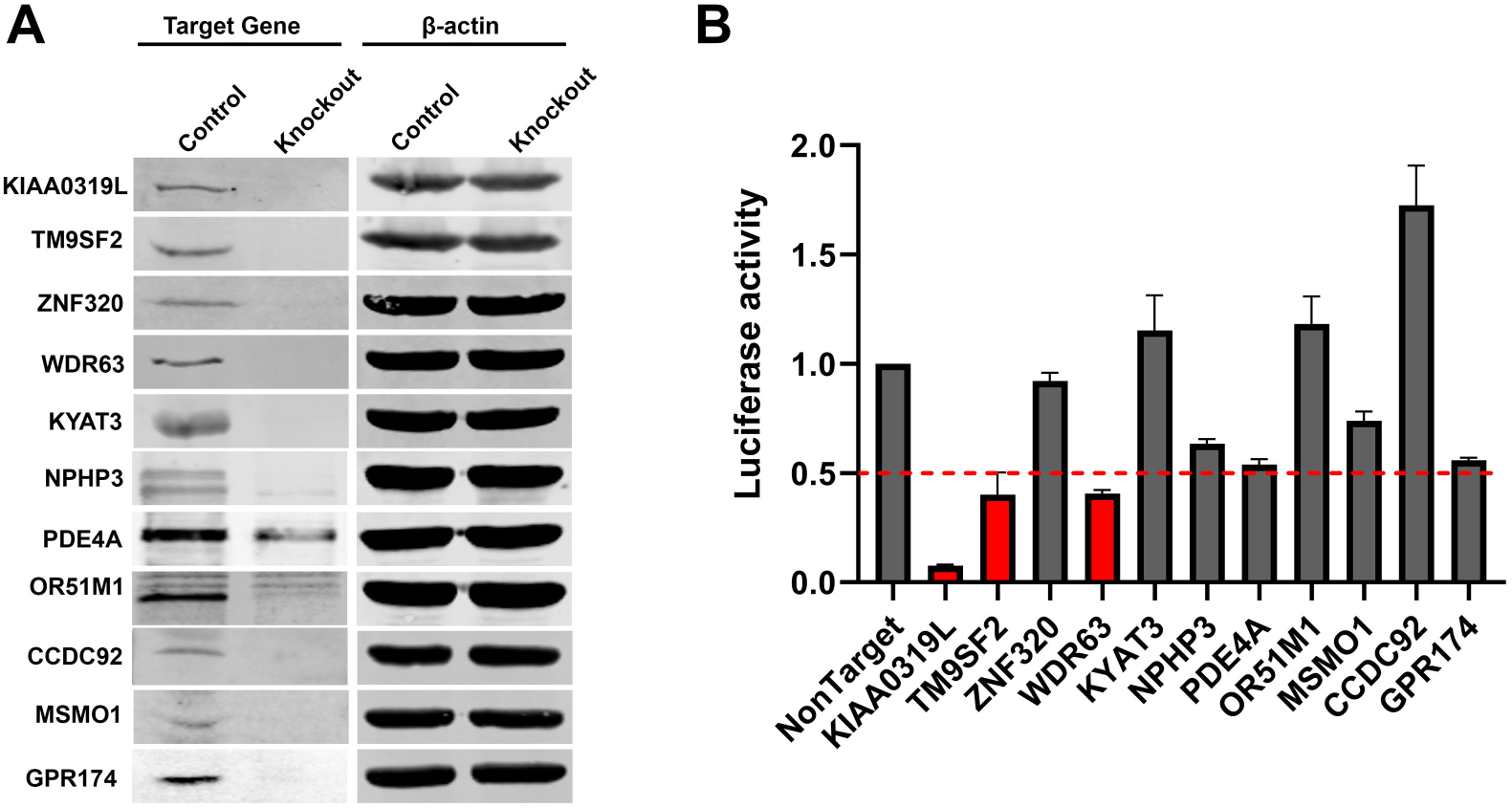
rAAV2.5 transduction of individual gene knockout HeLa cells. 11 candidate genes (*KIAA0319L*, *TM9SF2*, *ZNF320*, *KYAT3*, *NPHP3*, *WDR63*, *PDE4A*, OR51M1, *GPR174*, *CCDC92*, and *MSMO1*) were selected based on the shRNA knockdown screen. gRNA expressing lentiviral vector was applied in HeLa cells, and single-cell cloning was carried out to generate a knockout cell line. **(A) Western blotting.** Western blots show the knockout efficiency at 3 days post-transduction. β-actin was used as a loading control. **(B) Luciferase activities in gene knockout HeLa cells.** Gene knockout cells were transduced with rAAV2.5T at an MOI of 20,000 DRP/cell. At 3 days post-transduction, the luciferase activities were measured. The red dashed line indicates 50% of the luciferase activity in non-target control HeLa cells. Data shown are means with an SD from three replicates.

Collectively, we confirmed that *KIAA0319L*, *TM9SF2*, and *WDR63* are the top three host genes important for transduction and that *PDE4A* and *GPR174* also play a significant role in rAAV2.5T transduction in HeLa cells.

### *KIAA0319L* and *WDR63* knockout significantly decreases AAV2.5T transduction in HAE-ALI cultures

Since rAAV2.5T has a high tropism to human airway epithelia (15,18), we investigated the function of *KIAA0319L* and *WDR63* in rAAV2.5T transduction of polarized human airway epithelium cultured at an air-liquid interface (HAE-ALI). To this end, we used gRNA-expressing lentivirus to knock out *KIAA0319L* and *WDR63*, respectively, in an immortalized human airway epithelial cell line, CuFi-8. CuFi-8 cells maintain a phenotype of airway basal cells in proliferating culture and can undergo differentiation into pseudostratified HAE when cultured at an ALI (22) (**Figure 4A**). After confirmation of the gene knockout in proliferating CuFi-8 cells by Western blotting and genomic DNA sequencing (**Figure 4B**), we seeded the gene knockout CuFi-8 cells on collagen-coated transwells to differentiate them into HAE-ALI cultures. The maturation of epithelial differentiation is indicated by a value > 1,000 Ω.cm^2^ of the transepithelial electrical resistance (TEER). After 3 weeks of differentiation at ALI, the TEER of the HAE-ALI^ΔKIAA0319L^ and HAE-ALI^ΔWDR63^ cultures, as well as the non-target (NT) control cultures were measured. The TEER readings of these cultures approached ∼2,000 Ω.cm^2^, indicating the HAE-ALI cultures were fully differentiated (**Figure 4C**). We also examined the epithelial tight junction formation and cilial development by immunofluorescent staining for the tight junction protein ZO-1 and the cilial development marker β-tubulin IV. As shown in **Figure 4D**, both *KIAA0319L* and *WDR63* knockout did not obviously change the structure of ZO1 and cilia. Thus, we successfully developed HAE-ALI cultures with *KIAA0319L* or *WDR63* gene knockouts, with retained tight junction and cilia structures, as well as the normal epithelial barrier function.

**Figure 4.**
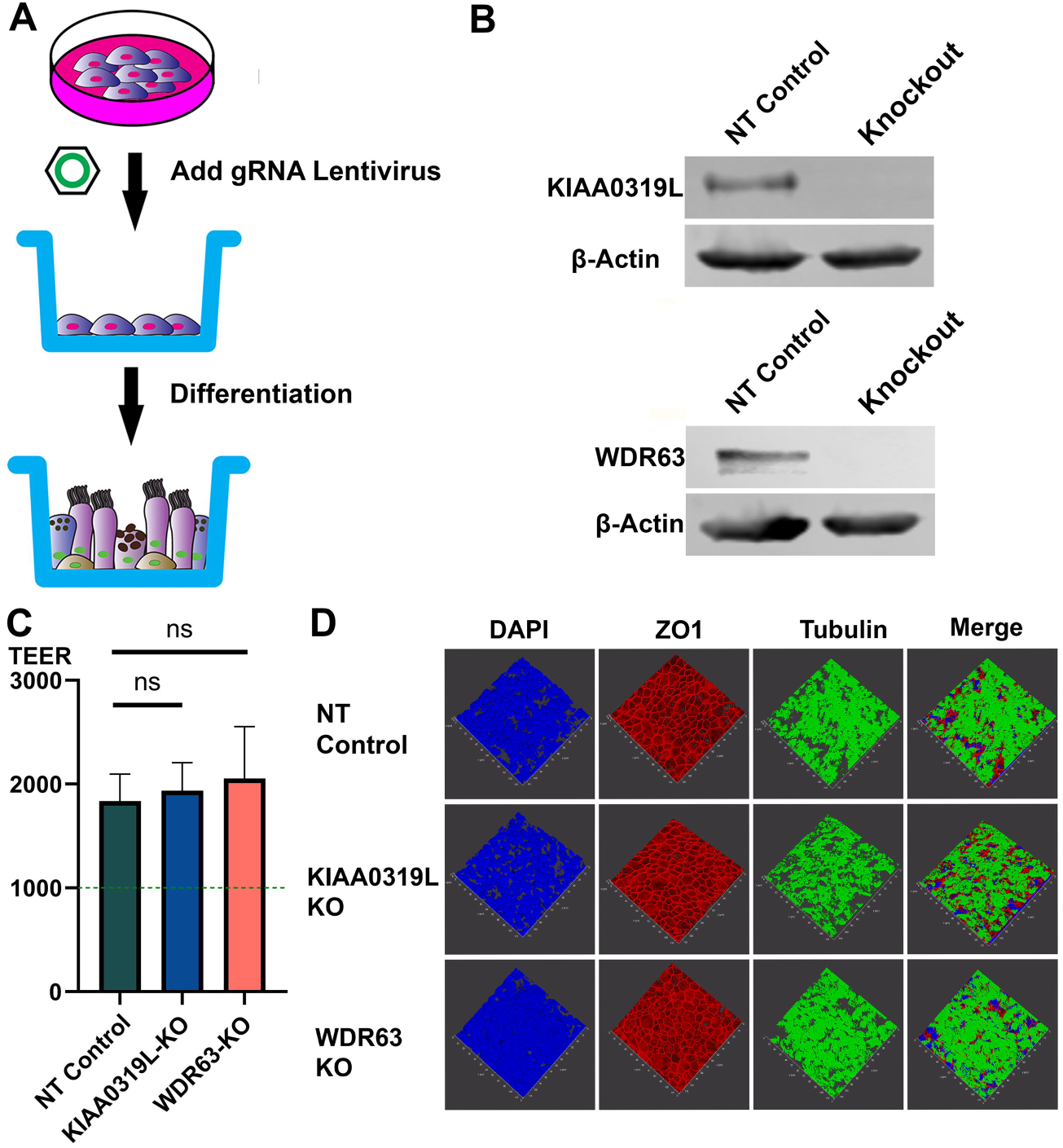
Generation of gene knockout in HAE-ALI cultures. **(A) Knockout of *KIAA0319L* and *WDR63* in HAE-ALI cultures.** A schematic diagram shows the generation of gene knockout of CuFi-8-derived HAE-ALI cultures. CuFi-8 cells were cultured in collagen-coated 100 mm dishes until confluent, then gRNA-expressing lentiviruses were added into the dishes. Under the selection of puromycin, the CuFi-8 cells were single-clone expanded for the production of a gene knockout cell line. Expanded gene knockout cells were transferred onto the transwell for polarization at ALI. **(B) Gene knockdown efficiency.** Western blotting and genomic DNA sequencing validated the knockout of *KIAA0319L* and *WDR63* in HAE-ALI cultures. Cells harvested from the transwells were extracted for proteins and DNA, respectively. Western blotting detected both the non-target control and knockout groups, KIAA0319L and WDR63, respectively. β-actin was detected as a loading control. The genomic DNA was amplified for sequencing the mutations at the gRNA targeting sequences of *KIAA0319L* and *WDR63* genes, respectively. The gene sequences show the original gene sequence, the non-target gene sequence, and the gene knockout sequence. **(C) TEER measurement.** After 1-month differentiation at ALI, HAE-ALI cultures, non-target NT (NT) control, KIAA0319L-KO, and WDR63-KO, were detected for TEER values. Data shown are means with an SD of three replicates. **(D) Three-dimensional confocal imaging of ZO-1 and** β**-tubulin expression in gene knockout HAE-ALI.** NT, KIAA0319L-KO, and WDR63-KO HAE-ALI cultures were fixed and co-stained with anti-β-tubulin IV (green) and anti-ZO-1 (red) antibodies. Nuclei were stained with DAPI (blue). A set of confocal images were taken at a magnification of × 40 (Leica SP8 STED) from the stained piece of the epithelium from the objective (z-axis) and reconstituted as a three-dimensional image as shown in each channel of fluorescence.

Next, the HAE-ALI^ΔKIAA0319L^ and HAE-ALI^ΔWDR63^ cultures were infected with rAAV2.5T at an MOI of 20,000 from the apical side (**Figure 5A**) or the basolateral side (**Figure 5B**). Infection of HAE-ALI^NT^ was included as the NT control. At 5 days post-transduction, the rAAV2.5T transduction efficiency was assessed by mCherry fluorescence and luciferase activity. HAE-ALI^ΔKIAA0319L^ and HAE-ALI^ΔWDR63^ showed a significant decrease in both mCherry and luciferase expression from the transduction at either the apical or basolateral side, compared to the NT control (**Figure 5A-D**). The luciferase quantification demonstrated that the knockout of KIAA0319L and WDR63 decreased the apical transduction by >70% and 71%, as well as basal transduction by >88% and 64%, respectively, compared to the NT control (**Figure 5C&D**). For all three types of HAE-ALI cultures, transduction at the apical side was higher than at the basolateral side.

**Figure 5.**
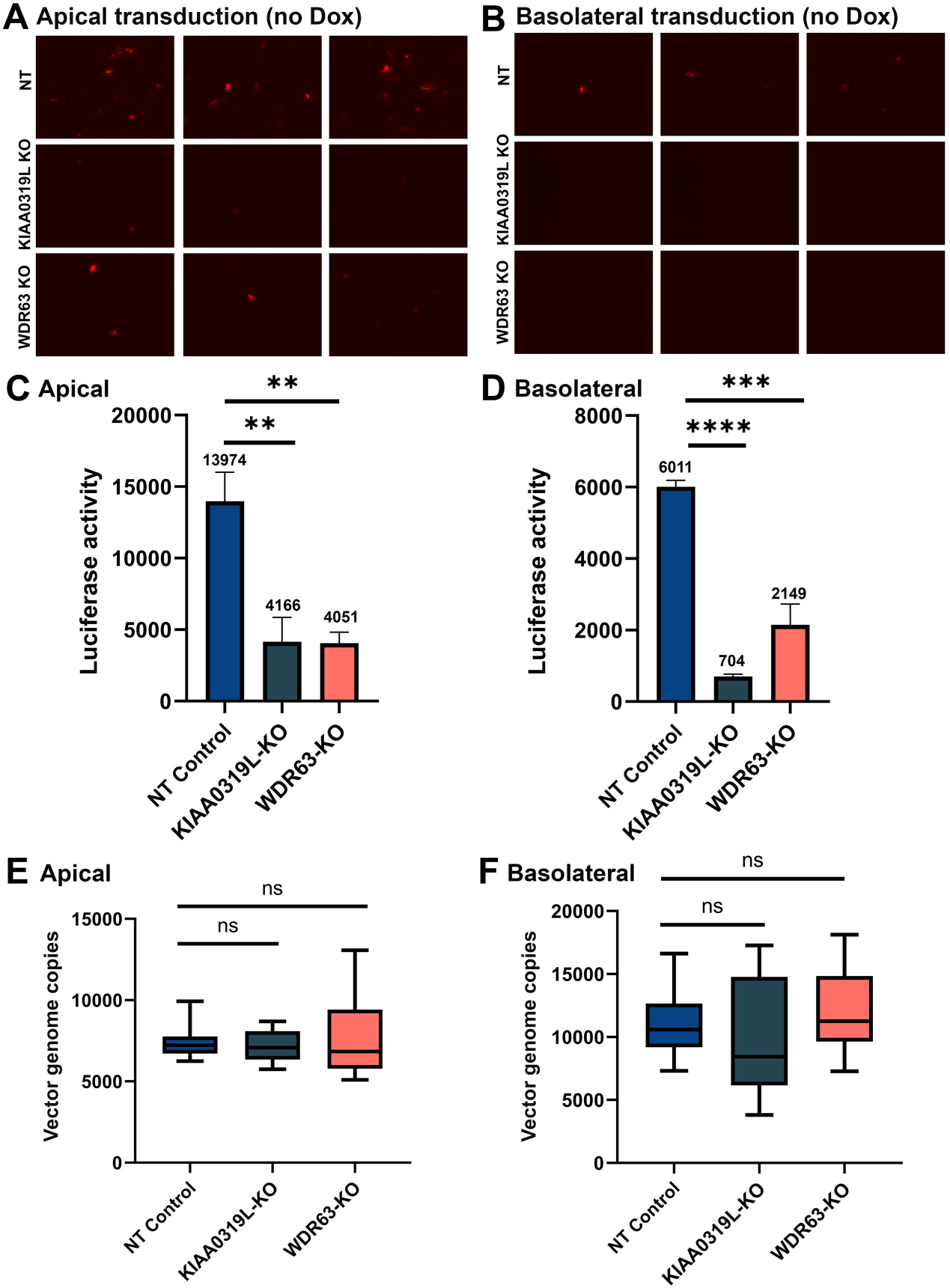
Knockout of KIAA0319L and WDR63 causes a significant decrease in transduction efficiency of rAAV2.5T in HAE-ALI cultures but not vector internalization. HAE-ALI cultures were transduced with rAAV2.5T at an MOI of 20,000 DRP/cell from the apical (left panels A, C and E) and basolateral chambers (right panels B, D, and F), respectively. After 16 hours post-transduction, both the apical and basolateral chambers were refreshed with culture media. **(A&B) mCherry expression.** Images were taken under a fluorescent imager at 5 days post-transduction. **(C&D) Luciferase activity assay.** Luciferase activity was measured at 5 days post-transduction. Data shown were means with an SD from three replicates**. (E&F) Vector internalization assay.** HAE-ALI cultures were incubated with rAAV2.5T at an MOI of 20,000 DRP/cell from the apical and basolateral chambers, respectively, for two hours at 37°C. Then the viruses were removed and both chambers were treated with Accutase three times. Viral DNA was extracted using the Pathogen viral DNA collection kit (Zymo), and quantified via qPCR using a mCherry gene-targeting probe to detect the entered viral genomes. The boundary of the box closest to zero indicates the 25^th^ percentile, a black line within the box marks the median, and the boundary of the box farthest from zero indicates the 75^th^ percentile. Whiskers above and below the box indicate the 10^th^ and 90^th^ percentiles.

As KIAA0319L was reported to function as a multi-serotype AAV receptor (12–14), we carried out a vector internalization assay to examine the role of KIAA0319L and WDR63 in vector endocytosis. The internalization assay showed that both HAE-ALI^ΔKIAA0319L^ and HAE-ALI^ΔWDR63^ did not have a significant decrease in the levels of internalized vectors, compared to the HAE-ALI^NT^ control from transduction either at the apical or basolateral side (**Figure 5E&F**). Differently, internalization assays in HeLa cells showed that *KIAA0319L* KO HeLa cells showed a three-fold decrease in vector internalization, compared with the control HeLa cells (**Figure S4**). Therefore, our virus internalization argues against the role of KIAA0319L in rAAV2.5T entry of human airway epithelia.

Doxorubicin (Dox), an FDA approved drug for cancer therapy, has been used as a potent agent to augment AAV-mediated transgene expression, especially in airway epithelia, functioning as a proteasome inhibitor to prevent the degradation of ubiquitinated AAV in the cytoplasm and facilitate vector nuclear import (23). We used rAAV2.5T to transduce HAE-ALI cultures in the presence of Dox at 2 µM during a 16-hour infection period. Dox treatment significantly increased mCherry expression from rAAV2.5T apical and basal transduction in HAE-ALI cultures at 5 days post-transduction (**Figure 6A&B**), compared to the no Dox treatment control (**Figure 5A&B**). From the luciferase activity, Dox treatment increased the rAAV2.5T transduction by 59- and 37-fold from the transduction of the apical and basolateral sides, respectively (**Figure 6C&D**). Luciferase activity was decreased in HAE-ALI^ΔKIAA0319L^ by 93% and 95%, respectively, from transduction of apical and basolateral sides, and a 77% and 81% decrease in HAE-ALI^ΔWDR63^ (**Figure 6C&D**). Importantly, in Dox treated HAE-ALI, we also found that both HAE-ALI^ΔKIAA0319L^ and HAE-ALI^ΔWDR63^ did not show any significant decrease in vector internalization during either apical or basolateral transduction at 37°C for 2 hours (**Figure 6E&F**).

**Figure 6.**
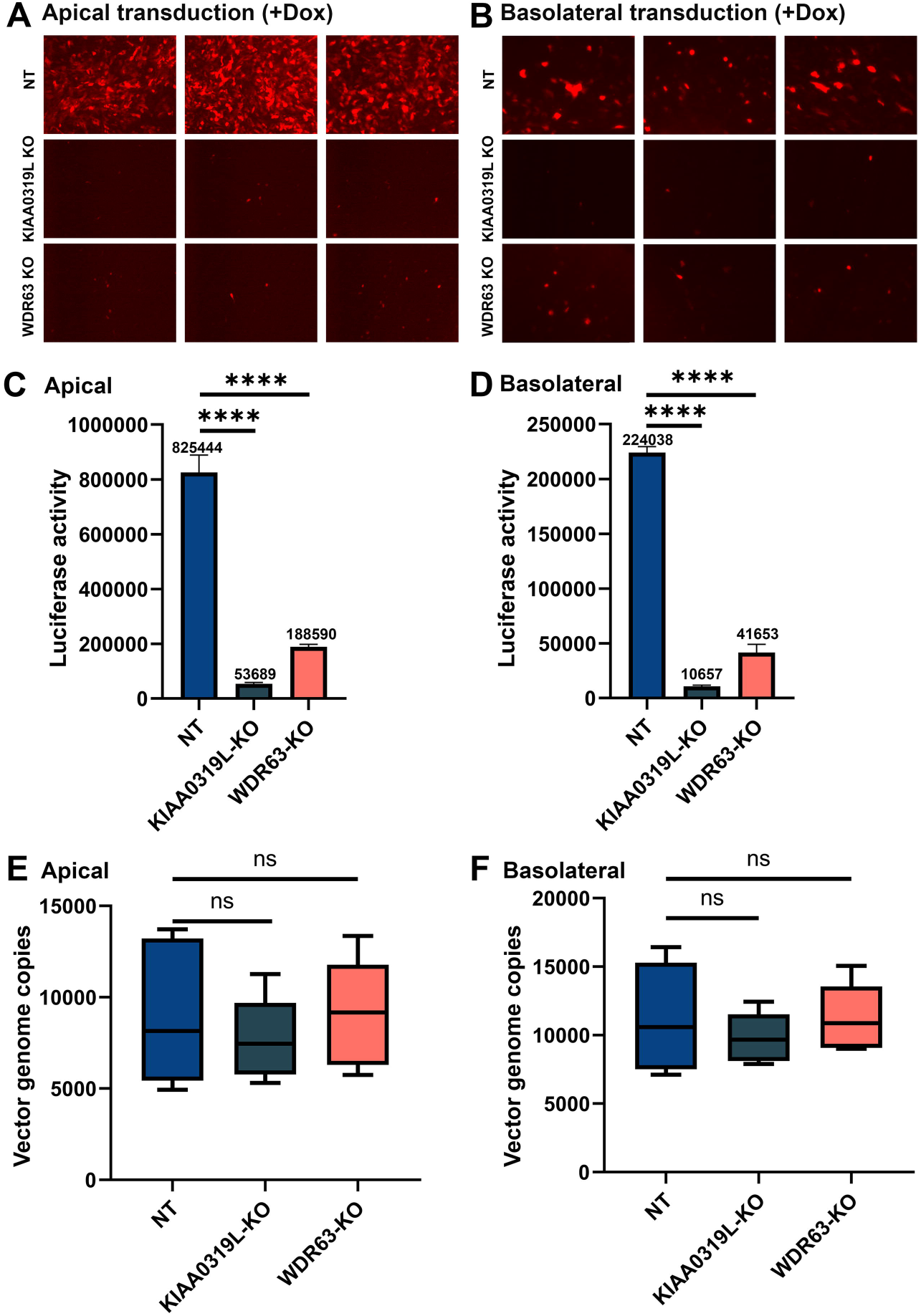
Knockout of *KIAA0319L* and *WDR63* causes a significant decrease in transduction efficiency of rAAV2.5T in Dox-treated HAE-ALI cultures but not vector internalization. HAE-ALI cultures were transduced with rAAV2.5T at an MOI of 20,000 DRP/cell from the apical (left panels A, C and E) and basolateral chambers (right panels B, D, and F), respectively. Dox was added with rAAV2.5T at 2.0 µM. After 16 hours post-transduction, both the apical and basolateral chambers were refreshed with culture media. **(A&B) mCherry expression.** Images were taken under a fluorescent imager at 5 days post-transduction. **(C&D) Luciferase activity assay.** Luciferase activity was measured at 5 days post-transduction. Data shown were means with an SD from three replicates**. (E&F) Vector internalization assay.** HAE-ALI cultures were transduced with rAAV2.5T from the apical and basolateral chambers, respectively, for two hours at 37°C. Then the vectors were removed and both chambers were treated with Accutase three times. Viral DNA was extracted and quantified via qPCR. The boundary of the box closest to zero indicates the 25th percentile, a black line within the box marks the median, and the boundary of the box farthest from zero indicates the 75th percentile. Whiskers above and below the box indicate the 10th and 90th percentiles.

Taken all these results together, although *KIAA0319L* and *WDR63* knockout drastically reduced rAAV2.5T transduction in HAE-ALI cultures, lack of KIAA0319L and WDR63 expression did not affect vector internalization into the cells during either apical or basolateral transduction. Notably, the treatment of Dox did not influence the entry of rAAV2.5T in HAE-ALI from either the apical or basal side, consistent with the previous observations in the transduction of other AAV serotypes in HAE-ALI, with less efficiencies (23).

### Cellular localization of KIAA0319L and WDR63 in the major types of airway epithelial cells of HAE-ALI cultures

Human airways are lined by a variety of cell types (24). The surface layer is primarily populated with ciliated cells as well as goblet and club secretory cells, the intermediate cells are club cells that can be differentiated to ciliated or secretory cells, and secured to the basement membrane are basal stem cells. Thus, we investigated the transduction efficiency of rAAV2.5T in the four major epithelial cell types: ciliated, basal, club, and goblet cells. Using flow cytometry of the cells dissociated from the inserts, we first determined the CuFi-8-derived HAE-ALI cultures were composed of ciliated, basal, club, and goblet cells at percentages of approximately 50%, 24%, 22%, and 4%, respectively (**Figure 7A**). Next, we analyzed rAAV2.5T transduced HAE-ALI cultures and found there were ∼35.4% mCherry-positive cells in the transduced cultures. Then, mCherry positive cells were sorted and analyzed by flow cytometry for the percentages of the four cell types using cell-type specific antibodies. We determined 45%, 19.5%, 34%, and 1.5% of the ciliated, basal, club, and goblet cells, respectively, were presented in the rAAV2.5T transduced (mCherry positive cell population) (**Figure 7B**). Representative immunofluorescent images showed that all four types of airway epithelial cells were transduced by rAAV2.5T but at various levels (**Figure 7C**).

**Figure 7.**
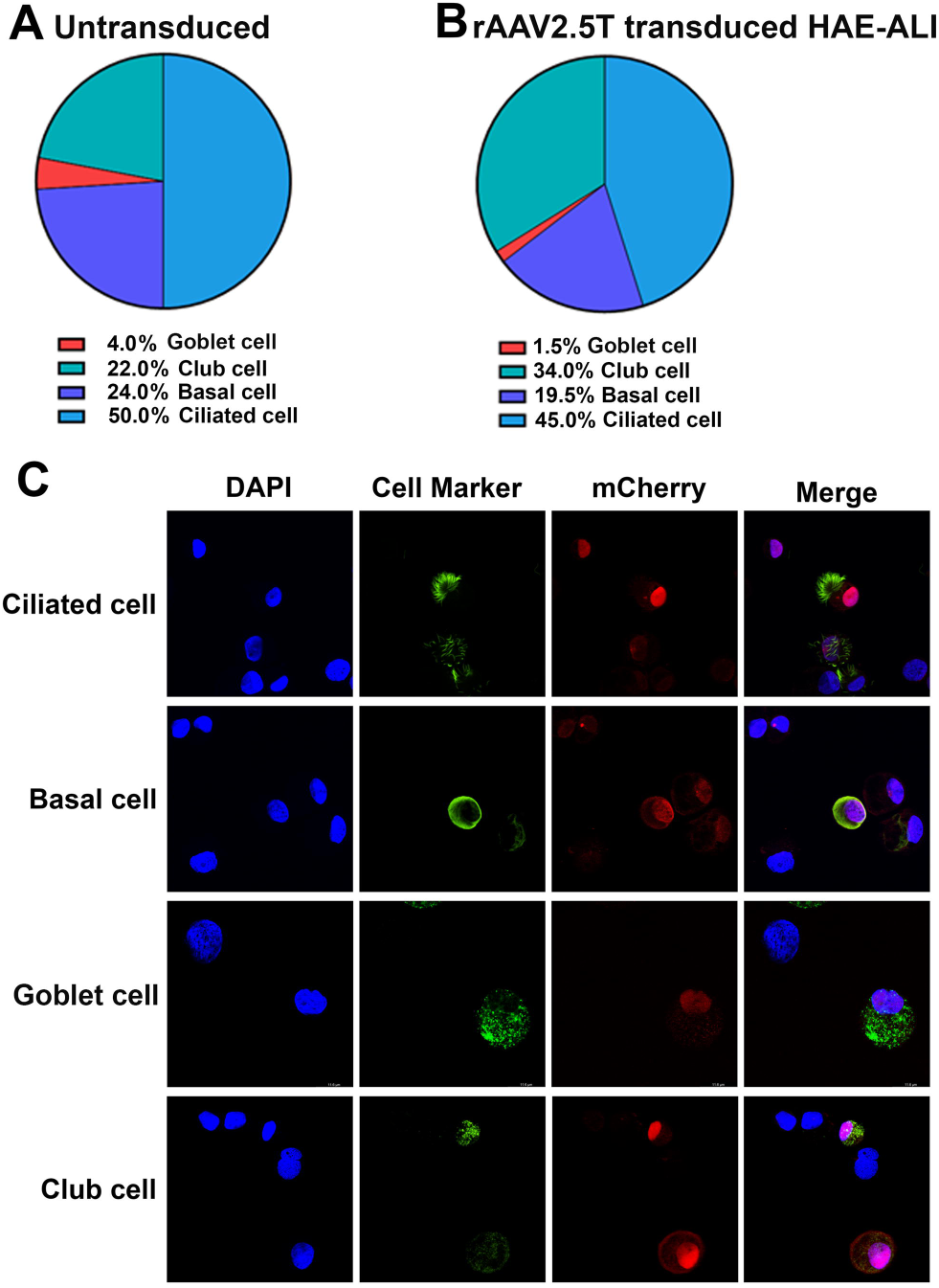
Analysis of rAAV2.5T transduced cell types in HAE-ALI cultures. HAE-ALI cultures were transduced with rAAV2.5T at an MOI of 20,000 DRP/cell from the apical chamber or mock transduced. Dox was added with rAAV2.5T at 2.0 µM. (**A**) **Flow cytometry of airway cell types in HAE-ALI cultures.** The cells of the mock transduced HAE-ALI cultures were digested off the transwell membranes, fixed and permeabilized, followed by immunostaining with the first antibody against each cell marker and an Alexa 488-conjugated secondary antibody. The stained cells were subjected to flow cytometry, and the percentage of each cell type was calculated. (**B**) **Flow cytometry of airway cells transduced with rAAV2.5T**. At 7 days post-transduction, the cells of the rAAV2.5T transduced HAE-ALI cultures were digested off the transwell membranes and sorted on a BD FACSAria™ III Cell Sorter. mCherry positive cells were sorted by FACS, fixed and permeabilized, followed by immunostaining with the first antibody against each cell marker and an Alexa 488-conjugated secondary antibody. The stained cells were subjected to flow cytometry, and the percentage of each cell type was calculated. **(C) Immunofluorescence assays.** At 7 days post-transduction, the cells of the rAAV2.5T transduced HAE-ALI cultures were digested off the transwell membranes and cytospun to slides, fixed and permeabilized, followed by co-immunostaining with an antibody against each cell marker. The stained cells were subjected to imaging under a confocal microscope at a magnitude × 100 (CSU-W1 SoRa, Nikon).

Knowing the localization of KIAA0319L and WDR63 in cells is critical to understand their functions during rAAV2.5T transduction of HAE-ALI. We carried out immunofluorescent staining to reveal the cellular localization of KIAA0319L and WDR63. The results showed that KIAA0319L localized on both the cell surface and in the cytoplasm of the four types of epithelial cells of HAE-ALI treated with Dox and apically transduced with rAAV2.5T (**Figure 8A**), indicating that KIAA0319L is likely to participate in the intracellular trafficking of rAAV2.5T (25). Strikingly, WDR63 localized both in the nucleus and cytoplasm with a preference to the nucleus (**Figure 8B**).

**Figure 8.**
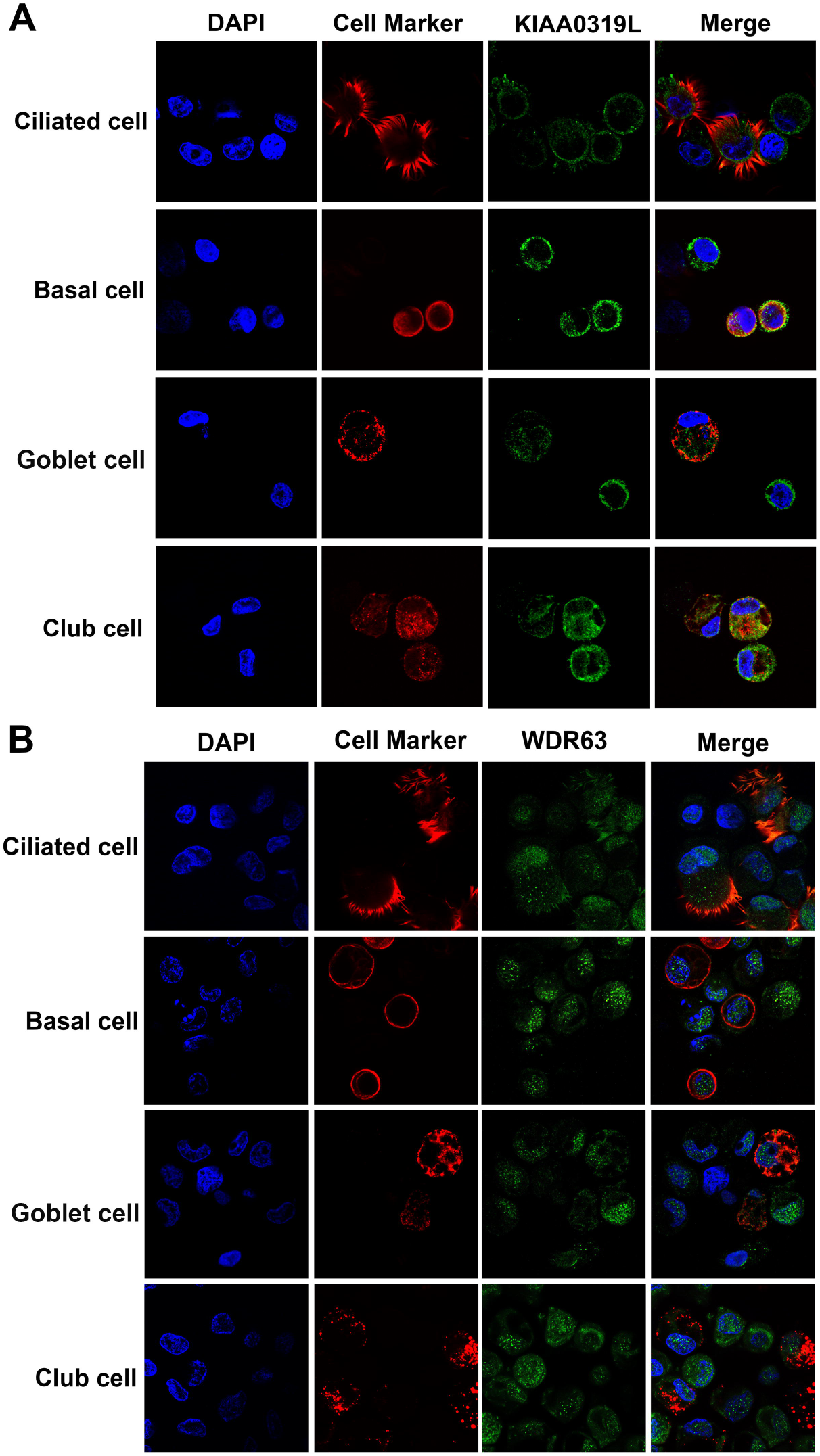
Localization of KIAA0319L and WDR63 in airway epithelial cells of HAE-ALI cultures. The cells of HAE-ALI cultures were digested off the transwell membranes with accutase, fixed and permeabilized, followed by co-immunostaining with an antibody against each cell marker and an anti-KIAA0319L (A) or anti-WDR63 (B), followed by staining of a correspondent secondary antibody. The stained cells were subjected to imaging under confocal microscopy at a magnitude × 100 (CSU-W1 SoRa, Nikon).

Lastly, we investigated rAAV2.5T capsid distribution in HAE-ALI cells using immunofluorescent assays. In ciliated cells, the nuclear entry of rAAV2.5T vectors was observed in nontarget control and HAE-ALI^ΔWDR63^ cultures, but vector particles were scarcely found within the nuclei of cells in HAE-ALI^ΔKIAA0319L^ cultures (**Figure 9A**). We also observed that no vector particles entered the nuclei of the club cells of the HAE-ALI^ΔKIAA0319L^, but more vector particles entered the nuclei of the cells of the nontarget control and HAE-ALI^ΔWDR63^ cultures (**Figure 9B**). We also fractionated the nucleus and cytosol of the three type HAE-ALI cultures and quantified the vector genome in both fractions. We found that vector genomes in the nuclei of HAE-ALI^ΔKIAA0319L^ were significantly decreased from 63.5% to 40.3%, but significantly increased in the nuclei of HAE-ALI^ΔWDR63^ from 63.5% to 70.5%, compared with the nontarget (NT) control (**Figure 9C**), suggesting that WDR63 may prevent the nuclear import of the vector. Notably, immunofluorescent assays found significant absence of the vector particles within the nuclei of the cells in transduced HAE-ALI^ΔKIAA0319L^, but substantial vial genome (40.3%) were still detectable in the nuclear fraction extracted from the transduced HAE-ALI^ΔKIAA0319L^. We reason this to the limitation of the fractionation method, as the results align with similar experiments carried out in a previously published study (26).

**Figure 9.**
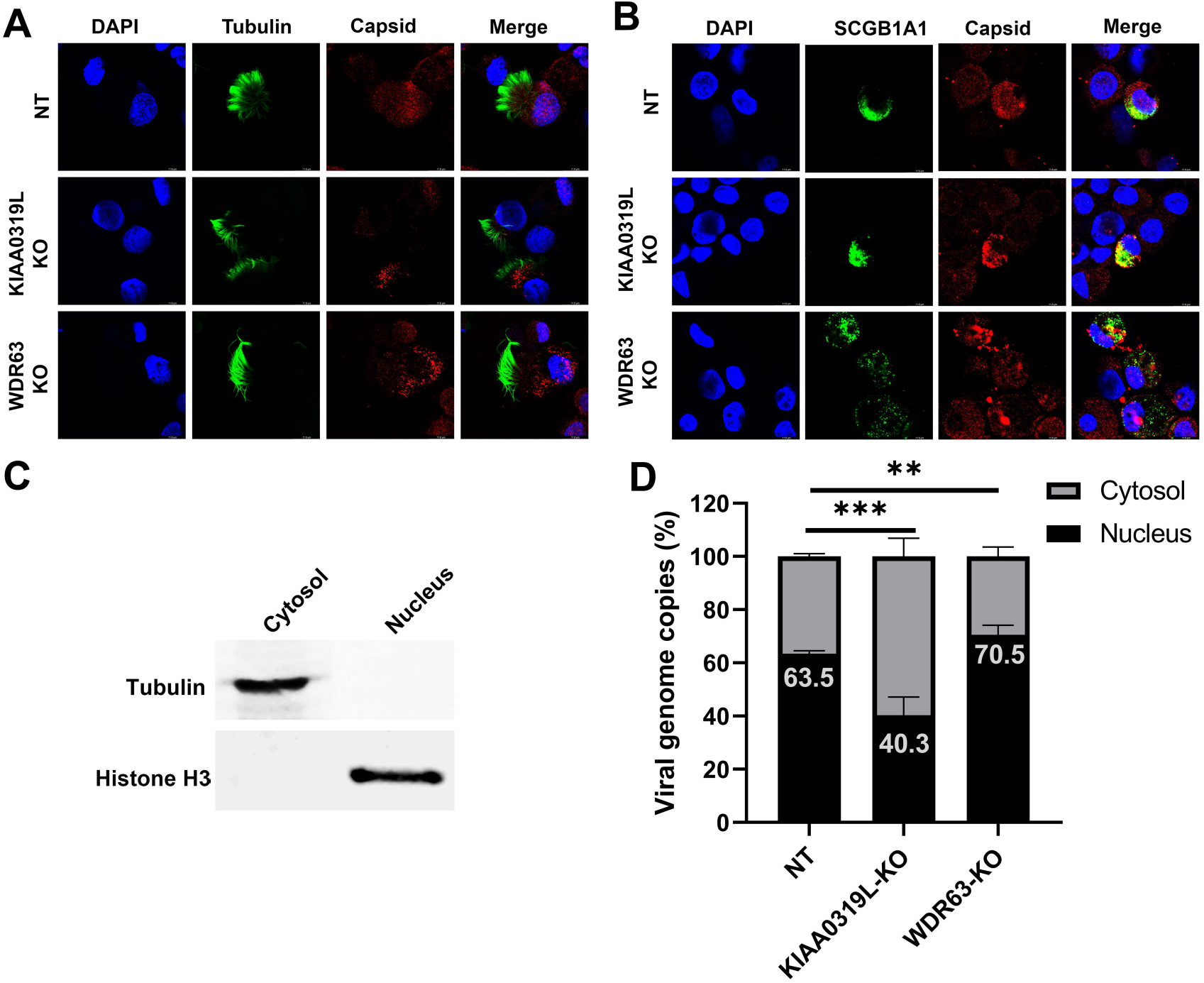
Localization of AAV2.5T capsid in transduced HAE-ALI cultures. AAV2.5T was used to apically transduce HAE-ALI cultures as indicated at an MOI of 20,000. **(A&B) rAAV2.5T capsid staining.** At 3 days post-transduction, cells were digested off the transwell membranes and cytospun onto slides. The slides were fixed and permeabilized, followed by co-immunostaining of ciliated cell marker (Tubulin, A) or club cell marker (SCGB1A1; B) with AAV2.5T capsid. The stained cells were subjected to imaging under confocal microscopy at a magnitude of × 100 (CSU-W1 SoRa, Nikon). **(C&D) rAAV2.5T genome distribution.** At 3 days post-transduction, cells in the transwell were washed with Accutase three times and then dissociated by incubation with Accutase for 1 hour. After washing, a cell fraction extraction kit (ThermoFisher) was used to extract the cytosol and nucleus. (C) Western blotting of the cytosol and nucleus fractions extracted from the nontarget control HAE-ALI cultures using anti-Tubulin and anti-Histone H3, respectively. (D) Quantification of viral genome in each fraction. Viral DNA was extracted from each fraction and quantitated using qPCR. Error bars represent the standard error of the mean (SEM) from three transwell replicates.

Taken together, our results suggest KIAA0319L and WDR63 are critical host factors for rAAV2.5T transduction in HAE-ALI cultures. KIAA0319L KO did not impact the internalization of rAAV2.5T vector but decreased the nuclear import of the vector in HAE-ALI. Thus, the critical role of KIAA0319L is confined to the intracellular trafficking of the vector to the nucleus. In contrast, WDR63 KO significantly increased the nuclear import of rAAV2.5T, suggesting that WDR63 may prevent nuclear import of the vector.

## Discussion

rAAVs enter cells and exhibit tissue tropism through a multistep process involving viral attachment, internalization, trafficking, and productive transduction (nuclear import, uncoating and transgene expression). The exact mechanisms can vary depending on the AAV serotype and the target cell type. While the tropism is considered broad in general, rAAVs inefficiently transduce lung airways from apical side. rAAV2.5T is an AAV variant developed from directed evolution for the treatment of the CF lung disease via gene therapy. It has demonstrated a higher efficiency in correcting the deficient Cl^-^ transportation in the HAE-ALI cultures derived from the lung donors of CF patient following apical transduction when compared to rAAV1 (27), which is the most efficient airway transduction vector from naturally occurring AAV serotypes (28). rAAV1 uses both α2,3 and α2,6 sialic acid residues that are present on N-linked glycoproteins as primary attachment receptors for transduction (29), but rAAV2.5T utilizes only α2,3 N-linked sialic acid residues, like AAV5 (18). While a previous study found the lack of endogenous KIAA0319L expression on the apical surface membrane of HAE-ALI, suggesting that the efficient rAAV2.5T transduction is accomplished with an KIAA0319L independent pathway (20), nothing is known about the mechanism underlying the highly efficient gene delivery of rAAV2.5T into human airway epithelia. In this study, we identified that KIAA0319L is a critical factor for rAAV2.5T transduction of airway epithelia, but not for vector internalization by acting as the proteinaceous cellular receptor. In addition, we confirmed WDR63, a less studied WD repeat domain containing protein, is an important host factor for rAAV2.5T transduction in human airway epithelia. Similar to KIAA0319L, WDR63 does not contribute to vector internalization. Importantly, we also identified that novel host factors, PDE4A and GPR174, play a moderate role in rAAV2.5T transduction in HeLa cells.

AAVs bind to proteoglycans as a primary attachment receptor, which facilitates the cellular uptake/internalization (30), for example, AAV2 binds to heparan sulfate proteoglycans (HSPG) (31,32). Post attachment, AAVs are thought to employ a proteinaceous receptor(s) to mediate cellular entry. The identification of proteinaceous receptors for AAV uptake has been a focus of AAV research for a long journey (33,34). A 150-kDa AAV-binding protein was initially revealed by Mizukami and Brown in 1996 on a virus overlay assay (35). In 2016, Carette and Chapman’s labs collaboratively identified KIAA0319L as a multi-serotype AAV receptor (AAVR) and demonstrated the physical interaction between KIAA0319L and AAV2 (12). KIAA0319L is both a N- and O-glycosylated protein with a size of 150 kDa (14), and the glycosylation of KIAA0319L is not essential for AAV2-AAVR interactions or AAV2 transduction. KIAA0319L is significantly involved in the transduction of AAV1-3 and 5-9; however, it is not essential for AAV4 transduction (13).

The capsid of AAV2.5T only has one mutation of A581T differed from AAV5 in the major capsid protein VP3; however, rAAV2.5T transduction exhibits a tropism to human airway epithelia much higher than its parent AAV5 (15). Molecular modeling of AAV2.5T showed the A581T mutation occurs at the mouth of the predicted AAV5 sialic acid binding pocket. An anti-KIAA0319L antibody blocked transduction of AAV2 from the basolateral side but not AAV2.5T from the apical side, suggesting that there is an unknown unique apical receptor for AAV2.5T (20). Since the anti-KIAA0319L polyclonal antibody was used at a single dose (50 µg/ml), it remains possible KIAA0319L plays a role in apical transduction of HAE. In this study, we used an sgRNA library and performed two rounds of screening in HeLa S3 cells to identify AAV2.5T receptors and host restriction factors as monolayer airway epithelial cells, e.g., CuFi-8, are less transducible. Our screen results showed that KIAA0319L is the top-ranking gene restricting rAAV2.5T transduction of HeLa cells. When knocked out in human airway epithelia, a nearly 95% reduction of rAAV2.5T transduction was observed from both apical and basolateral infection. However, in our study, the knockout of KIAA0319L did not significantly decrease vector internalization, regardless of whether a high or low MOI was used (data no shown), Interestingly, rAAV2.5T internalization decreased by ∼50% after knocking out KIAA0319L in HeLa cells, compared to WT HeLa cells, suggesting that the function of KIAA0319L might differ based on the cell type. We propose that KIAA0319L plays an important role in the intracellular trafficking of rAAV2.5T in airway epithelia (**Figure 10**). After internalization, AAV2.5T interacts with KIAA0319L in the endocytosed vesicle which then facilitates vector escape from the trans-Golgi network (TGN) or late endosome. This could be the physiological function of KIAA0319L, by binding to an unknown ligand and recycling from the cell surface and endosome to the perinuclear TGN. Thus, our study suggests there are other cell surface proteins that are required for rAAV2.5T internalization of airway epithelia, which have not been selected in the CRISPR screen in HeLa cells (**Figure 10**).

**Figure 10.**
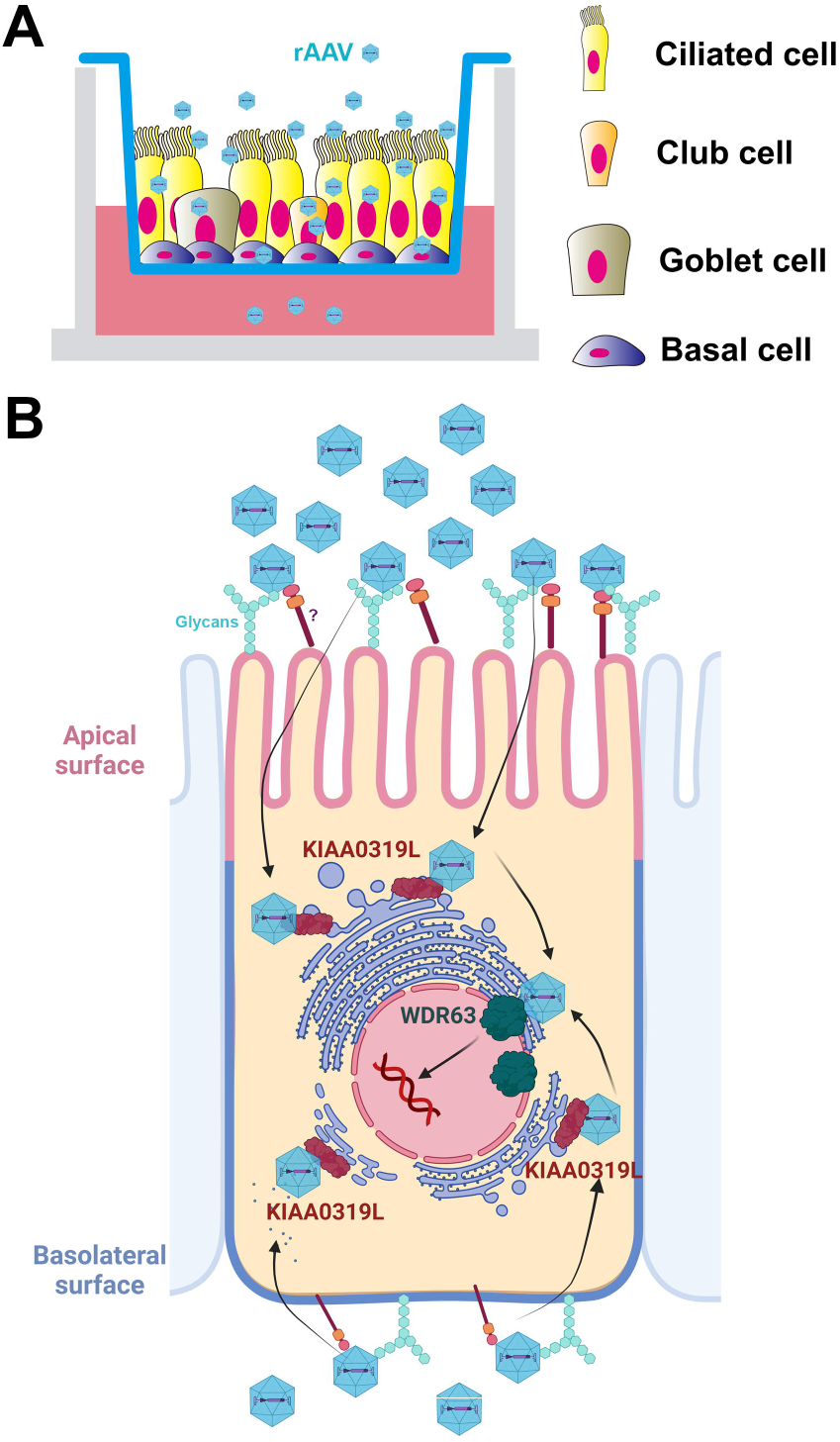
A model of rAAV2.5 transduction of HAE-ALI. (**A**) **Well-differentiated HAE-ALI model:** Human airway epithelial cells are differentiated on Transwell inserts at an air-liquid interface (ALI) for 3-4 weeks. Four major types of epithelial cells in the well differentiated polarized HAE-ALI cultures: basal, ciliated, goblet, and club cells are diagrammed in the Transwell insert. rAAV2.5T transduces the major 4 types of airway epithelia cells. (**B**) **AAV2.5T entry of HAE.** Represented is a ciliated cell diagramed with the nucleus (N) surrounded with trans-Golgi network (TGN) and endoplasmic reticulum (ER). AAV2.5T binds glycans on and enter into airway epithelial cells of the HAE-ALI cultures from either apical or basolateral side through an unknown proteinaceous receptor (marked with “?”), independent of the KIAA0319L. The capsid likely interacts with KIAA0319L during entry and localizes to TGN, followed by nuclear input. WDR63 likely plays a role in transgene expression in the nucleus.

We identified novel host factors, WDR63, PDE4A, and GPR174, play a significant role in rAAV2.5T transduction in HeLa cells, among which WDR63 is confirmed to be critical for rAAV2.5T transduction in human airway epithelia. WDR63 is a WD repeat-containing protein that plays roles as a negative regulator of cell migration, invasion, and metastasis through inhibiting actin polymerization (36). WDR63 has also been shown as a positive enhancer for osteogenic stem cell differentiation and enhances the expression of *BSP*, *OSX*, and *RUNX2* (37), indicating that WDR63 might be a transcriptional factor to enhance transgene expression in the nucleus (**Figure 10B**). WDR63 is localized both in the nucleus and cytoplasm. We also found that WDR63 did not colocalize with the vector in cells (**Figure S5**). WDR63 is not required for vector internalization, but plays a negative role in AAV2.5T nuclear entry. The involvement of WDR63 in rAAV transduction of HAE-ALI was also validated using rAAV2 and rAAV5 (**Figure S6**), thereby confirming that the role of WDR63 in rAAV transduction is independent of the VP1u and is likely serotype independent as well. Additional efforts are apparently required to verify the role of PDE4A and GPR174 in the rAAV2.5T transduction in human airway epithelia.

HAE cultures derived from primary bronchial epithelial cells isolated from the airways of human lung donors or patients undergoing lung transplantation have been extensively used to study the biology of respiratory epithelium (38). However, limited by the difficulty of gene knockout in primary cells with limited proliferating capacity, the HAE-ALI culture we used were differentiated/polarized from CuFi-8 cells, which are immortalized human airway epithelial cells that were isolated from a cystic fibrosis patient and express hTERT and HPV E6/E7 genes (22). CuFi-8 derived HAE is composed of 50% ciliated cells, 24% basal cells, 22% club cells, and 4% goblet cells, which have a similar composition to the results of single-cell RNA sequencing of nasal epithelial culture (39). In rAAV2.5T transduced HAE at an MOI of 20,000 DRP/cell, about one-third (35.4%) of the airway epithelial cells expressed mCherry, supporting the high transduction efficiency of airway cells. Among all the mCherry positive (transduced) cells, club cells took 34% and basal cells took ∼20%. Both club cells and basal cells are progenitors of the airway epithelium (38,40). Importantly, basal and club cells can be differentiated into ciliated and goblet cells under certain conditions, such as pathogenic viral infection (41,42). For cystic fibrosis gene therapy, it is more beneficial to transduce the stem-like cells rather than the ciliated cells (43).

AAV has emerged as a prominent viral vector platform for gene therapy, holding great promise in the treatment of genetic diseases and cancers. However, underlying its biological processes remain largely unexplored. Gaining a comprehensive understanding of the mechanistic basis of rAAV transduction biology will enable us to take advantage of its full potential. The human airway epithelia are composed of multiple cell types, including basal stem cells. AAV transduction of various airway epithelial cells may use different cellular receptors and pathways, which is important for airway gene delivery using AAV vectors. Notably, the requirement of KIAA0319L for highly efficient airway epithelial gene delivery, regardless of basal or apical infection, but not for vector internalization highlights the unique feature of rAAV2.5T-mediated airway gene delivery.

## Materials and Methods

### Cells and cell culture

#### Cells

Both HEK293FT (ThermoFisher) and HeLa cells (ATCC CCL-2) were grown in Dulbecco’s Modified Eagle Medium (DMEM; HyClone #SH30022.01, Cytiva, New York, NY) supplemented with 10% FBS and 100 units/mL penicillin-streptomycin in a humidified incubator with 5% CO2 at 37°C. HeLa S3 cells (ATCC CCL-2.2) were grown in Ham F-12 (#10-080-CV, Corning. Corning, NY) supplemented with 10% fetal bovine serum (FBS; # F0926; MilliporeSigma, St. Louis, MO) and 100 units/mL penicillin-streptomycin. Cells were cultured in shaking flasks on an orbital shaker platform at 120 rpm in a humidified incubator with 5% CO_2_ at 37°C. The cells were maintained at a low density of 0.1-0.3 million/ml. CuFi-8 cells were immortalized from human primary airway epithelial cells isolated from a cystic fibrosis patient by expressing hTERT and HPV E6/E7 genes (22). They were cultured on collagen-coated 100-mm dishes in PneumaCult™Ex Plus medium (#05040; StemCell, Vancouver, BC,).

#### Human airway epithelium cultured at an air-liquid interface (HAE-ALI)

Proliferating CuFi-8 cells dissociated from flasks were directly loaded on collagen-coated Transwell permeable supports (#3470; Costar, Corning) with PneumaCult™Ex Plus in both apical and basal chambers. One day after seeding, media were replaced with PneumaCult™-ALI medium (# 05001; StemCell) for 1–2 days, then, the media in the apical chamber were removed. The cells were then differentiated/polarized in PneumaCult™-ALI medium at an air-liquid interface (ALI) for 3–4 weeks (44). The maturation of the polarized HAE-ALI cultures derived from CuFi-8 cells was determined by transepithelial electrical resistance (TEER) measured with a Millicell ERS-2 volt-ohm meter (MilliporeSigma, Burlington, MA). ALI cultures with a TEER value of >1,000 Ω·cm^2^ were used for experiments.

### Production of rAAV2.5T vectors

The dual report rAAV2.5T vector, rAAV2/2.5T.F5tg83luc-CMVmCherry, was produced using a triple-plasmid transfection method using calcium phosphate (28). Briefly, the AAV2 Rep and AAV5 capsid-expressing plasmid pRep2Cap2.5T (15), the adenovirus 5 helper pAd4.1 (45), and the AAV2 transgene plasmid pAV2F5tg83luc-CMVmCherry (46) were transfected at a molar ratio of 1:1:1 into HEK293 cells cultured in 150-mm plates flasks. rAAV2.5T vectors were purified in cesium chloride density gradients, following a method as previously described (28). Vector titers were determined by real-time quantitative polymerase chain reaction (qPCR) using primers and probes specific to the *mCherry* transgene as DNase I resistant particle (DRP) per ml (46).

### Genome-wide CRISPR screen

HeLa S3 cells were transduced with a Streptococcus pyogenes (sp)Cas9 expressing lentivirus (#52962; Addgene, Watertown, MA) at an MOI of 20 transduction units (TU)/cell. After 3 days post-transduction, Blasticidin S, hereafter referred to as “blasticidin” (ThermoFisher, Waltham, MA) was added at 10 μg/ml to select spCas9-expressing cells. Western blotting and immunofluorescence staining were performed to confirm spCas9 expression. After the cells were expanded to 100 million, human CRISPR Brunello lentiviral pooled libraries (#73179-LV; Addgene) were added to the cell culture at an MOI of 0.25 in the presence of polybrene (Santa Cruz, Dallas, Texas). At 24 hours post-transduction, puromycin was added at a concentration of 2 μg/mL to select sgRNA-expressing cells for 1 week to select the blasticidin and puromycin double-resistant (gene edited) cells for proliferation. 100 million cells were harvested for extraction of genomic DNA (gDNA) as the unselected control (gDNA^Ctrl^). Another 100 million cells were transduced with rAAV2.5T at an MOI of 20,000 DRP/cell, ensuring that over 95% of HeLa S3 cells were transduced. At 3 days post-transduction, the cells were subjected to sorting on a BD FACSAria™ III Cell Sorter to select the top 1% mCherry-negative cells at the Flow Cytometry Core of University of Kansas Medical Center. ∼1 million cells of mCherry-negative were collected and expanded to ∼100 million, which underwent the second round of rAAV2.5T transduction and sorting process. Again, ∼1 million mCherry-negative cells were collected and expanded to ∼100 million cells, and were subjected to extraction of gDNA, which was designated as gDNA^Screen^.

### Genomic DNA extraction, next generation sequencing (NGS), and bioinformatics analysis

#### Genomic DNA (gDNA) extraction

gDNA samples from the unselected control (gDNA^Ctrl^) and 2^nd^ (gDNA^Screen^) selected group cells were extracted using the Blood and Cell Culture DNA Midi Kit (#13343; QIAGEN, Germantown, MD).

#### NGS and bioinformatics analysis

The extracted gDNA samples were subjected to PCR-based amplification of guide sequences and indexed according to the protocol from the Broad Institute of MIT and Harvard (47). The PCR amplicons were sequenced with Illumina NextSeq 2000 platform. All the reads of sgRNAs were analyzed using the MAGeCK software package (48). Significance values were determined after normalization to the control population, and the data were reported as an enrichment score.

The analyzed data were visualized through GraphPad. For visualization purposes, genes were sorted by gene ontology terms. The hits [by −log_10_(enrichment score)] were then plotted along the y-axis and were arbitrarily scattered within their categories along the x-axis. The size of the dot was determined according to the selected/unselected fold change.

### Western blotting

Cells were dissociated from the flask/transwell and solubilized with RIPA buffer [150 mM NaCl, 1% NP-40, 0.5% sodium deoxycholate, 0.1% sodium dodecyl sulfate (SDS), 50 mM Tris-HCl, pH 7.4]. The cell lysates were mixed with 5 × protein loading buffer and boiled for 10 minutes. Then, the samples were subjected to SDS-10% polyacrylamide gel electrophoresis, followed by transferring the separated proteins onto a nitrocellulose membrane. After blocking with non-fat milk for 1 hour, the membrane was incubated with the primary antibody overnight, followed by incubation with an infrared dye-conjugated IgG (H+L) secondary antibody for 1 hour. Finally, the membrane was imaged on a Li-Cor Odyssey imager (LI-COR Biosciences, Lincoln, NB).

### rAAV Transduction

For HeLa cells, they were seeded overnight in 48-well plates. rAAV2.5T was added into each well at an MOI of 20,000 DRP/cell. The transduction efficiency was analyzed using a firefly luciferase assay and mCherry intensity at 3 days post-transduction.

For apical transduction of HAE-ALI, 400 µL of PBS-diluted rAAV2.5T was added into the apical chamber of the transwell at an MOI of 20,000 DRP/cell. Then, 1 ml of culture media with or without 2 µM doxorubicin was added to the basolateral chamber. After 16 hours, all liquid in the apical and basolateral chambers was removed and washed with D-PBS, pH7.4 (Corning) three times. Fresh culture media were then added to the basolateral chamber.

For basolateral transduction of HAE-ALI, the media in the basolateral chamber were removed and replaced with 1 ml of culture media with or without 2 µM doxorubicin. rAAV2.5T was then directly added to the basolateral chamber at an MOI of 20,000 DRP/cell. The media in the basolateral chamber were then replaced.

### Luciferase assays

Firefly luciferase activity was detected using the Luciferase Assay System (#E4550; Promega, Madison, WI). At 3 days post-transduction for Hela cells or 5 days post-transduction for HAE-ALI cultures, cell culture media were removed, and cells were lysed in 200 µL per 48-well of Lysis Buffer (Promega), then frozen at −80C. After thawing, firefly luciferase expression was measured in relative light units on a multi-mode microplate reader using (Synergy H1, Agilent).

### mCherry quantification

mCherry expression was visualized under an inverted fluorescent microscope (NIKON Ti-S), and the intensity was quantified by image J.

### Gene silencing using shRNA-expressing lentiviruses

All of the shRNA expressing lentivirus (shRNA-lenti), generated from shRNA cloned pLKO.1-puro, were purchased from MilliporeSigma (**Table S2**). HeLa cells were seeded in a 24-well plate with 80% cell confluence. After the culture media were removed, shRNA-lenti was added into each well at an MOI of ∼10 TU/cell. The plate was then placed in a humidified incubator with 5% CO_2_ at 37°C for 2 hours, followed by addition of 500 μL of culture media. At 2 days post-transfection, puromycin was added at 2 μg/mL (#P9620, MillporeSigma) to each well to select gene silenced cells. Western blot assay was performed to confirm the knockdown efficiency.

### Generation of gene knockout stable cell lines

#### pLentiCRISPRv2 constructs

sgRNAs targeting genes, listed in **Table S3**, were respectively cloned in plentiCRISPRv2 vector (49). The lentiCRISPRv2 expressing a scramble sgRNA (5’-GTA TTA CTG ATA TTG GTG GG-3’) was used as a non-target (NT) control.

#### gRNA-based lentivirus production

Lentivirus was produced in HEK293FT cells by transient transfection using PEI Max and performed according to a previously published protocol (49). Briefly, individual sgRNA-containing lentiviruses were produced in HEK293FT cells seeded at 4 × 10^6^ cells per 150-mm dish one night before transfection. One hour prior to transfection, the media were changed with fresh media without antibiotics, followed by transfection of psPAX2, pMD2G, and gRNA sequence cloned lentiCRISPR v2 plasmid, at a 2:1:2 ratio. PEI max (#24765; Polysciences Inc.) was added at a mass ratio of 1:3 DNA to PEI max. The supernatant was collected at 48 hours post-transfection, clarified by centrifugation at 3,000 rpm for 30 minutes, and filtered through a 0.45-μm filter. The filtrated supernatant (lentivirus) was concentrated in a SureSpin 630 rotor in Sovall WX 80 (TheromoFisher) at 24,000 rpm for 2 hours at 4°C. The concentrated lentiviruses were resuspended in DMEM media and stored at - 80°C.

#### Generation of stable cell lines

HeLa or CuFi-8 cells were seeded at a density of 1 × 10^5^ cells per well in a 24-well plate. For CuFi-8 cells, the plates were collagen-coated. When the cells reached 80% confluence, lentivirus containing sgRNAs was added to each well at ∼5 MOI. At two days post-transduction, puromycin was added at 2 µg/ml to select for gene knockout cells. The selected cells were then digested and seeded into a 96-well plate at a density of 0.5 cells/well. Once sufficient confluence was reached, the cells in each well were reseeded into a 6-well plate, and Western blot analysis was performed to determine the knockout efficiency.

### Vector internalization assay

HAE-ALI cultures were infected with rAAV2.5T at an MOI of 20,000 from the apical or basolateral side. HeLa cells were infected at the same MOI. The cultures/cells were then incubated in a humidified incubator with 5% CO_2_ at 37°C for 2 hours before being washed with D-PBS three times. We carried out vector internalization assay using a previously published method (50). Cells were washed with Accutase (#AT104, Innovative Cell Technologies, Inc.) three times and then incubated with Accutase at 37°C for 1 hour. After washing with PBS three times, viral DNA was collected using Quick-DNA/RNA Pathogen Kits (#R1042, Zymo Research) and quantified by qPCR using mCherry primers/probe.

### Nuclear and cytoplasmic extraction

We fractionated the nucleus versus cytosol fraction of the HAE-ALI cultures using the NE-PER™ Nuclear and Cytoplasmic Extraction Reagents (#78833, ThemroFisher).

### Immunofluorescence confocal microscopy

For immunofluorescence of intact HAE-ALI culture, the membrane support of the insert was cut off and fixed in 4% paraformaldehyde in PBS at 4°C overnight. The fixed membrane was washed in D-PBS three times and then split into 8 pieces for whole-mount immunostaining. The fixed membrane pieces were permeabilized with 0.5% Triton X-100 in D-PBS for 15 min at room temperature and mounted on a slide. The slide was incubated with a primary antibody in D-PBS with 2% FBS for 1 h at 37°C, followed by washing 3 times, and was incubated with Alexa 488 and Alexa 594 secondary antibodies, followed by staining of the nuclei with DAPI (4’,6-diamidino-2-phenylindole).

For analysis of the AAV2.5T transduced cells of HAE-ALI cultures, we dissociated the cells off the supportive membranes of the Transwell inserts by incubation with Accutase. After incubation for 1 h at 37°C, cells were completely detached from the membrane and well separated. The cells were then cytocentrifuged onto slides at 1,500 rpm for 3 min in a Shandon Cytospin 3 cytocentrifuge. The slides were fixed with 4% paraformaldehyde in D-PBS for 10 minutes and were washed with D-PBS for 5 min three times. The fixed cells were permeabilized with 0.5% Triton X-100 for 15 min at room temperature. Then, the slide was incubated with one or two primary antibodies (**Table S4**) in D-PBS with 2% FBS for 1 h at 37°C. After washing three times, the slide was incubated with Alexa 488 and/or Alexa 594 conjugated secondary antibodies, followed by staining of the nuclei with DAPI.

The slides were visualized under a Leica TCS SP8 or Nikon CSU-W1 SoRa confocal microscope at the Confocal Core Facility of the University of Kansas Medical Center. Images were processed with the Leica Application Suite X software.

### Flow cytometry

#### Sorting of mCherry positive cells of HAE-ALI

HAE-ALI cultures were pretreated with 2 μM doxorubicin and transduced with AAV2.5T at an MOI of 20,000 DRP/cell. After 7 days post-transduction, the cells on the insert membrane were associated using Accutase for 1 hour and spun down at 300 × g for 3 minutes. The mCherry-positive cells were sorted using a FACSAriaIII sorter.

#### Airway epithelial cell typing

Cells of the HAE-ALI cultures were digested with Accutase for 1 hour. After spinning at 300 g for 3 minutes, the cells were fixed with 4% PFA and permeabilized with 0.5% Triton X-100 in PBS. The first antibody for each cell marker was added to the cells at a 1:1,000 dilution and incubated for 1 hour. The cells were then washed with PBS three times, and the correspondent second antibody conjugated with Alexa 488 was added to the cells, followed by incubation for 1 hour. After three washes with PBS, the samples were analyzed by flow cytometry to quantify Alexa 488 fluorescence. Negative control cells were incubated with 3% bovine serum albumin (BSA) and the second antibody.

### Statistical analysis

All data were determined with the means and standard deviations (SD) obtained from at least three independent experiments by using GraphPad. Error bars represent means and SD. Statistical significance (P value) was determined by using unpaired (Student) t-test for comparison of two groups. ****, P<0.0001; ***, P<0.001; **, P<0.01; and *, P<0.05 were considered as statistically significant and n.s. represents statistically no significance.

## Data availability

All data used to evaluate the conclusions in this study are presented in the paper and/or the **Supplementary Materials**. NGS data are available from the National Center for Biotechnology Information Sequencing Read Archive (SRA) under accession numbers: SRR24507630 and SRR24507271 and BioProject under accession number: PRJNA971590.

## Acknowledgments

The study was supported by NIH grants AI150877, AI156448, and AI166293. We are indebted to Dr. Richard Hastings at the Flow Cytometry Core Laboratory of the University of Kansas Medical Center. We acknowledge the Flow Cytometry Core Laboratory, which is sponsored, in part, by the NIH/NIGMS COBRE grant P30 GM103326 and the NIH/NCI Cancer Center grant P30 CA168524. The super resolution confocal microscope Leica SP8 STED was supported by NIH S10 OD 023625 and the Nikon CSU-W1 SoRa was supported by NIH S10 OD 032207 at the University of Kansas Medical Center. The funder had no role in study design, data collection and interpretation, or the decision to submit the work for publication.

## References

1. Cotmore, S. F., M. Agbandje-McKenna, M. Canuti, J. A. Chiorini, A. M. Eis-Hubinger, J. Hughes, M. Mietzsch, S. Modha, M. Ogliastro, J. J. Penzes, D. J. Pintel, J. Qiu, M. Soderlund-Venermo, P. Tattersall, P. Tijssen, and Ictv Report Consortium. 2019. ICTV Virus Taxonomy Profile: Parvoviridae. J Gen.Virol. 100:367–368.

2. Wang, D., P. W. L. Tai, and G. Gao. 2019. Adeno-associated virus vector as a platform for gene therapy delivery. Nat.Rev.Drug Discov. 18:358–378.

3. Kuzmin, D. A., M. V. Shutova, N. R. Johnston, O. P. Smith, V. V. Fedorin, Y. S. Kukushkin, J. C. M. van der Loo, and E. C. Johnstone. 2021. The clinical landscape for AAV gene therapies. Nat Rev.Drug Discov. 20:173–174.

4. Bennett, J., J. Wellman, K. A. Marshall, S. McCague, M. Ashtari, J. DiStefano-Pappas, O. U. Elci, D. C. Chung, J. Sun, J. F. Wright, D. R. Cross, P. Aravand, L. L. Cyckowski, J. L. Bennicelli, F. Mingozzi, A. Auricchio, E. A. Pierce, J. Ruggiero, B. P. Leroy, F. Simonelli, K. A. High, and A. M. Maguire. 2016. Safety and durability of effect of contralateral-eye administration of AAV2 gene therapy in patients with childhood-onset blindness caused by RPE65 mutations: a follow-on phase 1 trial. Lancet. 388:661–672.

5. Mueller, C., G. Gernoux, A. M. Gruntman, F. Borel, E. P. Reeves, R. Calcedo, F. N. Rouhani, A. Yachnis, M. Humphries, M. Campbell-Thompson, L. Messina, J. D. Chulay, B. Trapnell, J. M. Wilson, N. G. McElvaney, and T. R. Flotte. 2017. 5 Year Expression and Neutrophil Defect Repair after Gene Therapy in Alpha-1 Antitrypsin Deficiency. Mol.Ther. 25:1387–1394.

6. Lovric, J., M. Mano, L. Zentilin, A. Eulalio, S. Zacchigna, and M. Giacca. 2012. Terminal differentiation of cardiac and skeletal myocytes induces permissivity to AAV transduction by relieving inhibition imposed by DNA damage response proteins. Mol.Ther. 20:2087–2097.

7. Mendell, J. R., S. A. Al-Zaidy, L. R. Rodino-Klapac, K. Goodspeed, S. J. Gray, C. N. Kay, S. L. Boye, S. E. Boye, L. A. George, S. Salabarria, M. Corti, B. J. Byrne, and J. P. Tremblay. 2021. Current Clinical Applications of In Vivo Gene Therapy with AAVs. Mol.Ther. 29:464–488.

8. George, L. A., P. E. Monahan, M. E. Eyster, S. K. Sullivan, M. V. Ragni, S. E. Croteau, J. E. J. Rasko, M. Recht, B. J. Samelson-Jones, A. MacDougall, K. Jaworski, R. Noble, M. Curran, K. Kuranda, F. Mingozzi, T. Chang, K. Z. Reape, X. M. Anguela, and K. A. High. 2021. Multiyear Factor VIII Expression after AAV Gene Transfer for Hemophilia A. N.Engl.J.Med. 385:1961–1973.

9. Riyad, J. M. and T. Weber. 2021. Intracellular trafficking of adeno-associated virus (AAV) vectors: challenges and future directions. Gene Ther. 28:683–696.

10. Samulski, R. J. and N. Muzyczka. 2014. AAV-Mediated Gene Therapy for Research and Therapeutic Purposes. Annu.Rev.Virol. 1:427–451.

11. Meyer, N. L. and M. S. Chapman. 2022. Adeno-associated virus (AAV) cell entry: structural insights. Trends Microbiol. 30:432–451.

12. Pillay, S., N. L. Meyer, A. S. Puschnik, O. Davulcu, J. Diep, Y. Ishikawa, L. T. Jae, J. E. Wosen, C. M. Nagamine, M. S. Chapman, and J. E. Carette. 2016. An essential receptor for adeno-associated virus infection. Nature. 530:108–112.

13. Dudek, A. M., S. Pillay, A. S. Puschnik, C. M. Nagamine, F. Cheng, J. Qiu, J. E. Carette, and L. H. Vandenberghe. 2018. An Alternate Route for Adeno-associated Virus (AAV) Entry Independent of AAV Receptor. J Virol. 92:e02213–17.

14. Pillay, S., W. Zou, F. Cheng, A. S. Puschnik, N. L. Meyer, S. S. Ganaie, X. Deng, J. E. Wosen, O. Davulcu, Z. Yan, J. F. Engelhardt, K. E. Brown, M. S. Chapman, J. Qiu, and J. E. Carette. 2017. AAV serotypes have distinctive interactions with domains of the cellular receptor AAVR. J.Virol. 91:e00391–17.

15. Excoffon, K. J., J. T. Koerber, D. D. Dickey, M. Murtha, S. Keshavjee, B. K. Kaspar, J. Zabner, and D. V. Schaffer. 2009. Directed evolution of adeno-associated virus to an infectious respiratory virus. Proc.Natl.Acad.Sci.U.S.A. 106:3865–3870.

16. Ostedgaard, L. S., T. Rokhlina, P. H. Karp, P. Lashmit, S. Afione, M. Schmidt, J. Zabner, M. F. Stinski, J. A. Chiorini, and M. J. Welsh. 2005. A shortened adeno-associated virus expression cassette for CFTR gene transfer to cystic fibrosis airway epithelia. Proc.Natl.Acad.Sci.U.S.A. 102:2952–2957.

17. Tang, Y., S. Fakhari, E. D. Huntemann, Z. Feng, P. Wu, W. Y. Feng, J. Lei, F. Yuan, K. J. Excoffon, K. Wang, M. P. Limberis, R. Kolbeck, Z. Yan, and J. F. Engelhardt. 2023. Immunosuppression reduces rAAV2.5T neutralizing antibodies that limit efficacy following repeat dosing to ferret lungs. Mol.Ther.Methods Clin.Dev. 29:70–80.

18. Dickey, D. D., K. J. Excoffon, J. T. Koerber, J. Bergen, B. Steines, J. Klesney-Tait, D. V. Schaffer, and J. Zabner. 2011. Enhanced sialic acid-dependent endocytosis explains the increased efficiency of infection of airway epithelia by a novel adeno-associated virus. J.Virol. 85:9023–9030.

19. Afione, S., M. A. DiMattia, S. Halder, P. G. Di, M. Agbandje-McKenna, and J. A. Chiorini. 2015. Identification and mutagenesis of the adeno-associated virus 5 sialic acid binding region. J Virol. 89:1660–1672.

20. Hamilton, B. A., X. Li, A. A. Pezzulo, M. H. Abou Alaiwa, and J. Zabner. 2019. Polarized AAVR expression determines infectivity by AAV gene therapy vectors. Gene Ther. 26:240–249.

21. Meisen, W. H., Z. B. Nejad, M. Hardy, H. Zhao, O. Oliverio, S. Wang, C. Hale, M. M. Ollmann, and P. J. Collins. 2020. Pooled Screens Identify GPR108 and TM9SF2 as Host Cell Factors Critical for AAV Transduction. Mol.Ther.Methods Clin.Dev. 17:601–611.

22. Zabner, J., P. Karp, M. Seiler, S. L. Phillips, C. J. Mitchell, M. Saavedra, M. Welsh, and A. J. Klingelhutz. 2003. Development of cystic fibrosis and noncystic fibrosis airway cell lines. Am.J.Physiol Lung Cell Mol.Physiol. 284:L844–L854.

23. Yan, Z., R. Zak, Y. Zhang, W. Ding, S. Godwin, K. Munson, R. Peluso, and J. F. Engelhardt. 2004. Distinct classes of proteasome-modulating agents cooperatively augment recombinant adeno-associated virus type 2 and type 5-mediated transduction from the apical surfaces of human airway epithelia. J.Virol. 78:2863–2874.

24. Davis, J. D. and T. P. Wypych. 2021. Cellular and functional heterogeneity of the airway epithelium. Mucosal.Immunol. 14:978–990.

25. Pillay, S. and J. E. Carette. 2017. Host determinants of adeno-associated viral vector entry. Curr.Opin.Virol. 24:124–131.

26. Dudek, A. M., N. Zabaleta, E. Zinn, S. Pillay, J. Zengel, C. Porter, J. S. Franceschini, R. Estelien, J. E. Carette, G. L. Zhou, and L. H. Vandenberghe. 2020. GPR108 Is a Highly Conserved AAV Entry Factor. Mol.Ther. 28:367–381.

27. Tang, Y. and Z. Yan. 2020. Repeat Dosing of AAV2.5T to Ferret Lungs Elicits an Antibody Response That Diminishes Transduction in an Age-Dependent Manner. Mol.Ther.Methods Clin.Dev. 19:186–200.

28. Yan, Z., D. C. Lei-Butters, N. W. Keiser, and J. F. Engelhardt. 2012. Distinct transduction difference between adeno-associated virus type 1 and type 6 vectors in human polarized airway epithelia. Gene Ther. 20:972–983.

29. Wu, Z., E. Miller, M. Agbandje-McKenna, and R. J. Samulski. 2006. Alpha2,3 and alpha2,6 N-linked sialic acids facilitate efficient binding and transduction by adeno-associated virus types 1 and 6. J.Virol. 80:9093–9103.

30. Madigan, V. J. and A. Asokan. 2016. Engineering AAV receptor footprints for gene therapy. Curr.Opin.Virol. 18:89–96.

31. Summerford, C. and R. J. Samulski. 1998. Membrane-associated heparan sulfate proteoglycan is a receptor for adeno-associated virus type 2 virions. J.Virol. 72:1438–1445.

32. Qiu, J., A. Handa, M. Kirby, and K. E. Brown. 2000. The interaction of heparin sulfate and adeno-associated virus 2. Virology. 269:137–147.

33. Qiu, J., H. Mizukami, and K. E. Brown. 1999. Adeno-associated virus 2 co-receptors? Nat.Med. 5:467–468.

34. Qiu, J. and K. E. Brown. 1999. Integrin alphaVbeta5 is not involved in adeno-associated virus type 2 (AAV2) infection. Virology. 264:436–440.

35. Mizukami, H., N. S. Young, and K. E. Brown. 1996. Adeno-associated virus type 2 binds to a 150-kilodalton cell membrane glycoprotein. Virology. 217:124–130.

36. Zhao, K., D. Wang, X. Zhao, C. Wang, Y. Gao, K. Liu, F. Wang, X. Wu, X. Wang, L. Sun, J. Zang, and Y. Mei. 2020. WDR63 inhibits Arp2/3-dependent actin polymerization and mediates the function of p53 in suppressing metastasis. EMBO Rep. 21:e49269.

37. Diao, S., D. M. Yang, R. Dong, L. P. Wang, J. S. Wang, J. Du, S. L. Wang, and Z. Fan. 2015. Enriched trimethylation of lysine 4 of histone H3 of WDR63 enhanced osteogenic differentiation potentials of stem cells from apical papilla. J.Endod. 41:205–211.

38. Fulcher, M. L., S. Gabriel, K. A. Burns, J. R. Yankaskas, and S. H. Randell. 2005. Well-differentiated human airway epithelial cell cultures. Methods Mol.Med. 107:183–206.

39. Ruiz, G. a. S., M. Deprez, K. Lebrigand, A. Cavard, A. Paquet, M. J. Arguel, V. Magnone, M. Truchi, I. Caballero, S. Leroy, C. H. Marquette, B. Marcet, P. Barbry, and L. E. Zaragosi. 2019. Novel dynamics of human mucociliary differentiation revealed by single-cell RNA sequencing of nasal epithelial cultures. Development. 146:dev177428.

40. Konkimalla, A., S. Konishi, Y. Kobayashi, M. P. Kadur Lakshminarasimha, L. Macadlo, A. Mukherjee, Z. Elmore, S. J. Kim, A. M. Pendergast, P. J. Lee, A. Asokan, L. Knudsen, J. J. Bravo-Cordero, A. Tata, and P. R. Tata. 2022. Multi-apical polarity of alveolar stem cells and their dynamics during lung development and regeneration. iScience. 25:105114.

41. Hao, S., K. Ning, C. A. Kuz, K. Vorhies, Z. Yan, and J. Qiu. 2020. Long-Term Modeling of SARS-CoV-2 Infection of In Vitro Cultured Polarized Human Airway Epithelium. MBio. 11:e02852–20.

42. El-Badrawy, M. K., N. M. Shalabi, M. A. Mohamed, A. Ragab, and H. W. Abdelwahab. 2016. Stem Cells and Lung Regeneration. Int.J.Stem Cells. 9:31–35.

43. Yan, Z., P. B. McCray, Jr., and J. F. Engelhardt. 2019. Advances In Gene Therapy For Cystic Fibrosis Lung Disease. Hum.Mol.Genet. 28:R88–R94.

44. Yan, Z., X. Deng, and J. Qiu. 2020. Human Bocavirus 1 Infection of Well-Differentiated Human Airway Epithelium. Curr.Protoc.Microbiol. 58:e107.

45. Yan, Z., N. W. Keiser, Y. Song, X. Deng, F. Cheng, J. Qiu, and J. F. Engelhardt. 2013. A novel chimeric adenoassociated virus 2/human bocavirus 1 parvovirus vector efficiently transduces human airway epithelia. Mol.Ther. 21:2181–2194.

46. Wang, Z., F. Cheng, J. F. Engelhardt, Z. Yan, and J. Qiu. 2018. Development of a Novel Recombinant Adeno-Associated Virus Production System Using Human Bocavirus 1 Helper Genes. Mol.Ther.Methods Clin.Dev. 11:40–51.

47. Sanson, K. R., R. E. Hanna, M. Hegde, K. F. Donovan, C. Strand, M. E. Sullender, E. W. Vaimberg, A. Goodale, D. E. Root, F. Piccioni, and J. G. Doench. 2018. Optimized libraries for CRISPR-Cas9 genetic screens with multiple modalities. Nat.Commun. 9:5416–07901.

48. Li, W., H. Xu, T. Xiao, L. Cong, M. I. Love, F. Zhang, R. A. Irizarry, J. S. Liu, M. Brown, and X. S. Liu. 2014. MAGeCK enables robust identification of essential genes from genome-scale CRISPR/Cas9 knockout screens. Genome Biol. 15:554–0554.

49. Ning, K., C. A. Kuz, F. Cheng, Z. Feng, Z. Yan, and J. Qiu. 2023. Adeno-Associated Virus Monoinfection Induces a DNA Damage Response and DNA Repair That Contributes to Viral DNA Replication. MBio. 14:e0352822.

50. Ning, K., W. Zou, P. Xu, F. Cheng, E. Y. Zhang, A. Zhang-Chen, S. Kleiboeker, and J. Qiu. 2023. Identification of AXL as a co-receptor for human parvovirus B19 infection of human erythroid progenitors. Sci.Adv. 9:eade0869.

