## Supplemental Figure S1-S6 and Table S2-4 for "Identification of Host Restriction Factors Critical for Recombinant AAV Transduction of Polarized Human Airway Epithelium"

### Supplemental Materials

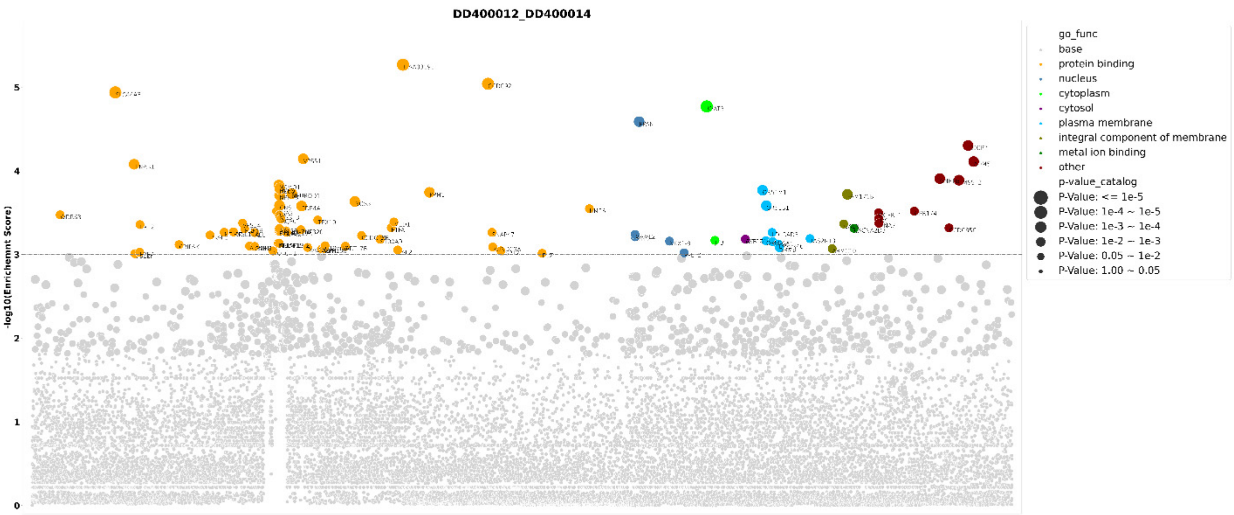


**Figure S1. Genes enriched in the second round of screen of mCherry negative cells.**

The x-axis shows all the genes enriched. The genes with an enrichment score of 3 are grouped by gene ontology analysis. Genes are indicated as circles with size corresponding to fold change of sgRNA reads in $gDNA^{Screen}$ and $gDNA^{Ctrl}$. The enrichment score [$-log_{10}$] of each gene based on MAGeCK analysis is shown in the y-axis.

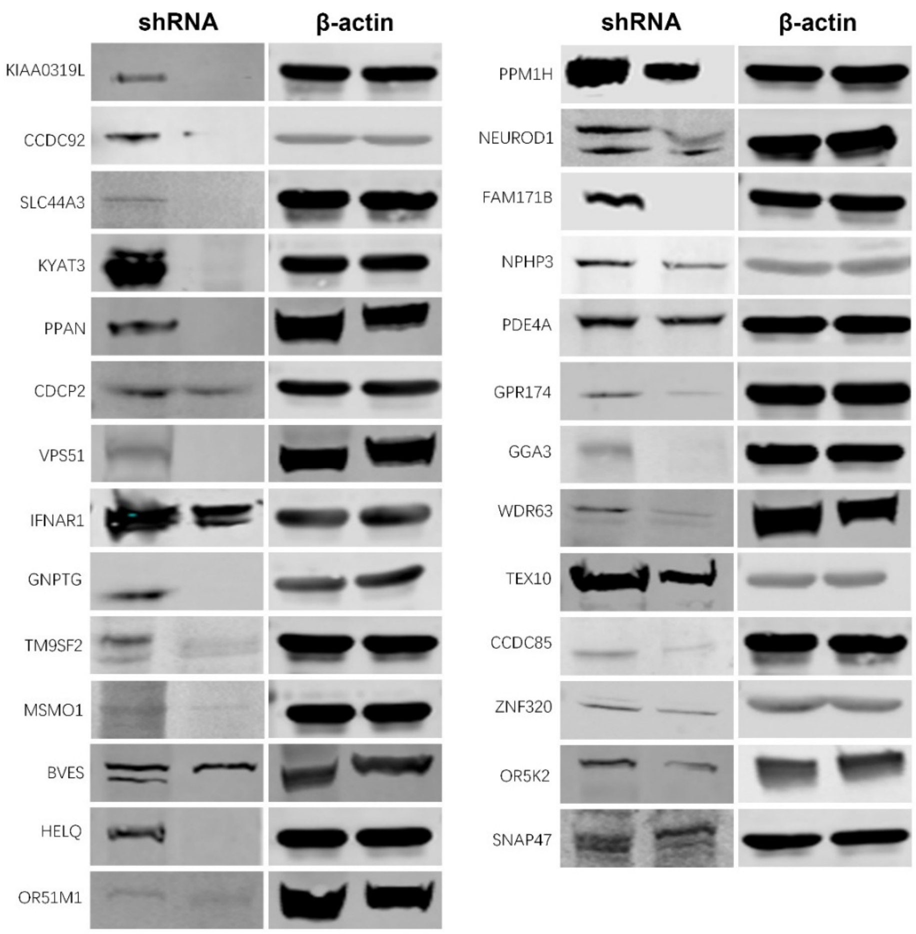


**Figure S2. Silencing of 27 candidate genes using shRNA-expressing lentiviruses in HeLa cells.**

HeLa cells seeded in 24-well plates were transduced with shRNA-expressing lentiviruses against the top 27 genes as indicated, respectively. After selection in puromycin, the cells were collected and lysed for Western blotting using antibodies against the proteins as indicated. β-actin was detected as a loading control.

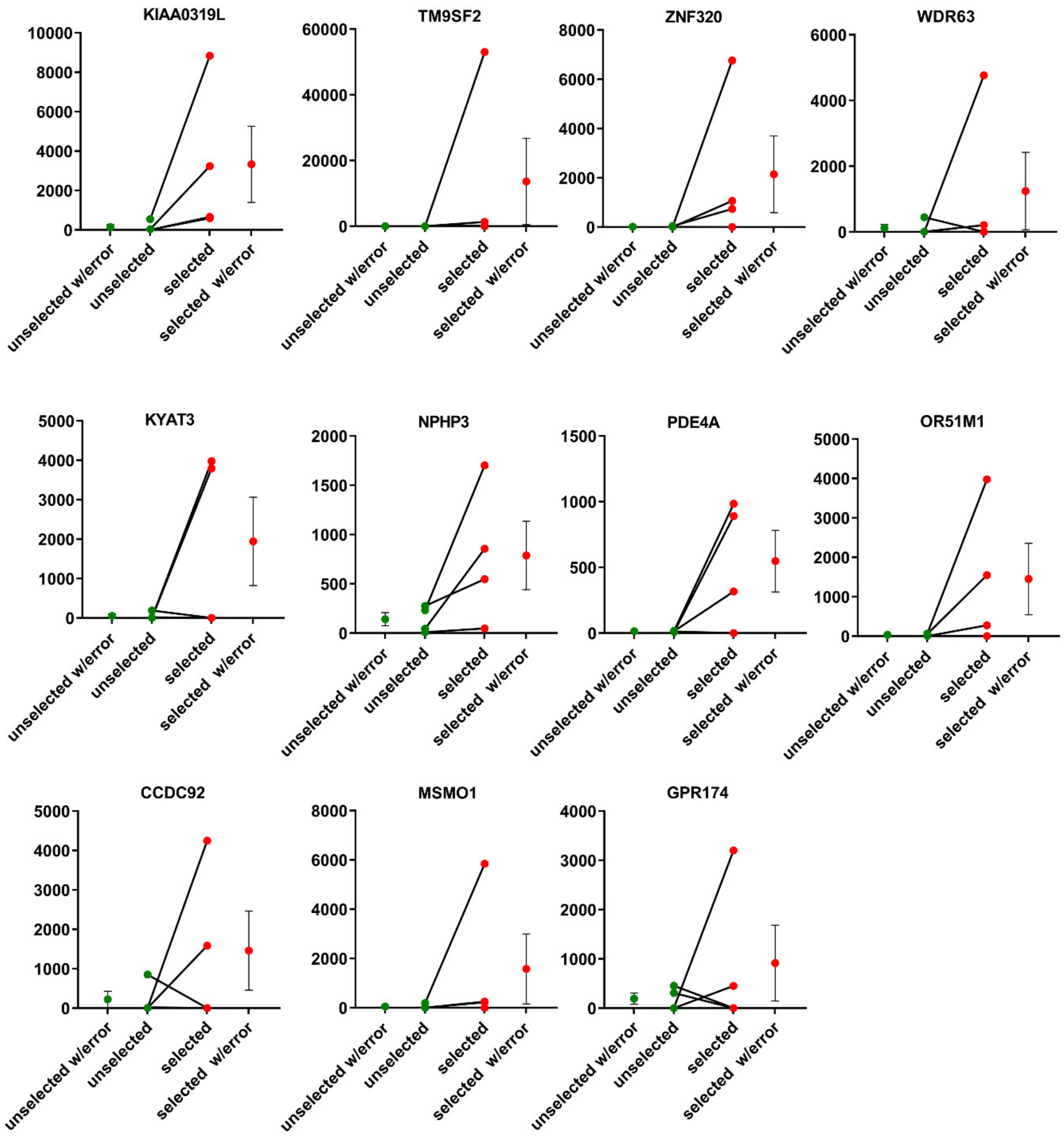


**Figure S3. Fold changes of the hits of the selected genes in the gRNA library screen using mCherry negative cells transduced with rAAV2.5T.**

This panel displays the individual gene fold-changes of 4 gRNAs per gene between the unselected and selected groups.

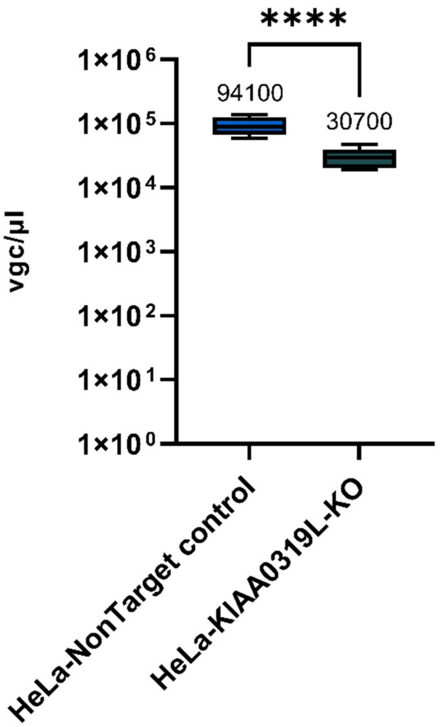


**Figure S4. rAAV2.5T internalization assay of HeLa cells.**

HeLa cells were incubated with rAAV2.5T at an MOI of 20,000 DRP/cell. After two hours at 37°C, the cells were used for vector internalization assay. The boundary of the box closest to zero indicates the 25th percentile, a black line within the box marks the median, and the boundary of the box farthest from zero indicates the 75th percentile. Whiskers above and below the box indicate the 10th and 90th percentiles.

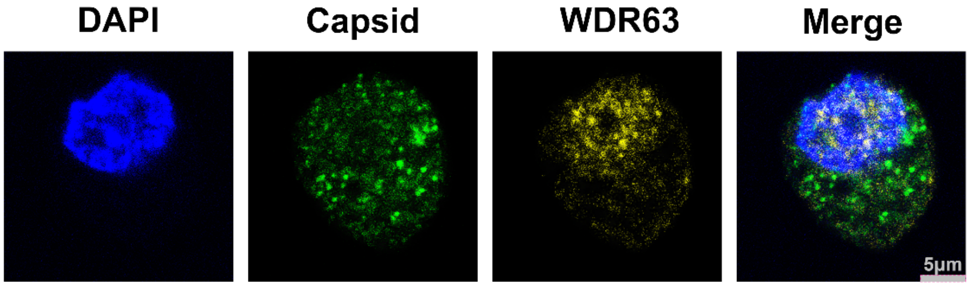


**Figure S5. Co-staining of AAV2.5T capsid with WDR63 in cells of HAE-ALI cultures.**

AAV2.5T was transduced into HAE-ALI cells at an MOI of 20,000. Three days after transduction, CuFi cells of the ALI culture were Accutase treated and cytospun onto slides for staining. The stained cells were subjected to imaging under a confocal microscope at a magnitude × 100 (CSU-W1 SoRa, Nikon). Green represents viral capsid, yellow represents WDR63, and blue represents DAPI staining.

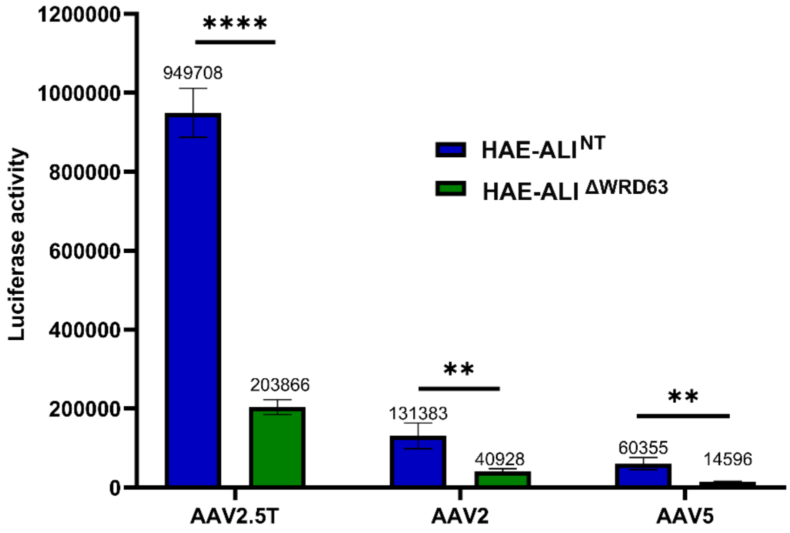


**Figure S6. Knockout of *WDR63* causes a significant decrease in transduction efficiency of rAAV2 and rAAV5 in Dox-treated HAE-ALI cultures.**

HAE-ALI cultures of non-target control (NT) or WRD63-knockout (KO) cells were transduced with rAAV2.5T, rAAV2, and rAAV5 that contain expression cassette of F5tg83gluc-CMVmCherry at an MOI of 20K from the apical side. Dox was added with rAAV2.5T at 2.0 μM. After 16 hours post-transduction, both the apical and basolateral chambers were refreshed with culture media. Luciferase activity was measured at 3 days post-transduction. Data shown were means with an SD from three replicates**.**

**Table S1. A list of genes enriched in the second round of mCherry negative cell screening and ranked by the -$\log_{10}$ enrichment score.**

NGS results of the sgRNA-targeted genes screened in the second-round screening ($gDNA^{Screen}$) were compared with the results from the background cells ($gDNA^{Ctrl}$) based on MAGeCK analysis. The analysis results are listed in **Supplemental Material 2**.

**Table S2. List of shRNAs used in the study.**

| *Gene* | **Cat. no. of shRNA at MilliporeSigma** |
|---|---|
| *Non-Target (NT) shRNA Control* | SHC016V |
| *KIAA0319L* | TRCN0000123109 |
| *CCDC92* | TRCN0000353704 |
| *SLC44A3* | TRCN0000158746 |
| *KYAT3* | TRCN0000150908 |
| *PPAN* | TRCN0000128301 |
| *CDCP2* | TRCN0000064638 |
| *VPS51* | TRCN0000151062 |
| *IFNAR1* | TRCN0000059013 |
| *GNPTG* | TRCN0000036049 |
| *TM9SF2* | TRCN0000059772 |
| *MSMO1* | TRCN0000230198 |
| *BVES* | TRCN0000153094 |
| *HELQ* | TRCN0000051559 |
| *OR51M1* | TRCN0000187514 |
| *PPM1H* | TRCN0000052772 |
| *NEUROD1* | TRCN0000019859 |
| *FAM171B* | TRCN0000020096 |
| *NPHP3* | TRCN0000118612 |
| *PDE4A* | TRCN0000048812 |
| *GPR174* | TRCN0000008261 |
| *GGA3* | TRCN0000232888 |
| *WDR63* | TRCN0000158457 |
| *TEX10* | TRCN0000364593 |
| *CCDC85C* | TRCN0000337094 |
| *ZNF320* | TRCN0000152095 |
| *OR5K2* | TRCN0000187322 |
| *SNAP47* | TRCN0000167217 |

**Table S3. gRNA sequences**

| ***Gene*** | **gRNA sequence (5’ → 3’)** |
|---|---|
| *KIAA0319L* | GAG GTG ACA CAA TAG CAA TG |
| *TM9SF2* | CTT GTT ACT TAT GTC CAT GG |
| *ZNF320* | GGA CAT CGT AGA GTT CAC AC |
| *WDR63* | TCC GGT TCA AGG AGA AAC AT |
| *KYAT3* | AAG GAC TTG ATA GTA ATG TG |
| *NPHP3* | TTA TTC ACT TAA CAT TAC CA |
| *PDE4A* | CAC AGT GCA CCA TGT TCC GG |
| *OR51M1* | GGC CTA ATT GTC ATC TTC CG |
| *CCDC92* | AGT GCT GGA GAA CAC CAT CA |
| *MSMO1* | AAG TTC CAG ATT GCA ACA TG |
| *GPR174* | TAT AAC CAT AGA ATA CCC AC |

**Table S4. The first antibodies used in this study.**

| Protein | Vendor | Cat. No. |
|---|---|---|
| KIAA0319L | Proteintech | 21016-1-AP |
| CCDC92 | Proteintech | 27192-1-AP |
| SLC44A3 | AB Clonal | A12820 |
| KYAT3 | Novus | 87387 |
| PPAN | AB Clonal | A7377 |
| CDCP2 | Novus | 87438 |
| VPS51 | AB Clonal | A15651 |
| IFNAR1 | AB Clonal | A1715 |
| GNPTG | Novus | 88443 |
| TM9SF2 | Novus | 95189 |
| MSMO1 | Novus | 59450 |
| BVES | AB Clonal | A0123 |
| HELQ | AB Clonal | A12661 |
| OR51M1 | Biorbyt | ORB398077 |
| PPM1H | Abcepta | AP9093 |
| NEUROD1 | Proteintech | 12081-1-AP |
| FAM171B | Novus | 93847 |
| NPHP3 | Proteintech | 22026-1-AP |
| PDE4A | Abcepta | AP17181 |
| GPR174 | Abcepta | AP16459 |
| GGA3 | Novus | HD120C2913 |
| WDR63 | Novus | 32639 |
| TEX10 | AB Clonal | A18118 |
| CCDC85C | Novus | 82622 |
| ZNF320 | Proteintech | 24882 |
| OR5K2 | Novus | 9805 |
| SNAP47 | Novus | 56894 |
| β-tubulin IV | MilliporeSigma | T7941 |
| ZO-1 | BD Bioscience | 610966 |
| Cytokeratin k5 | ThermoFisher Invitrogen | MA5-12596 |
| SCGB1A1 | ThermoFisher Invitrogen | MAB4218 |
| MUC5AC | Santa Cruz Biotechnology | sc-33667 |
| AAV5 Capsid | ARP | 608021 |
| β-Actin | MilliporeSigma | A5441 |
| Histon H3 | Proteintech | 17168-1-AP |
